# Genomic and metabolic characterization of *Trueperella pyogenes* isolated from domestic and wild animals

**DOI:** 10.1101/2024.09.09.612034

**Authors:** Gabriela Magossi, Katherine E. Gzyl, Devin B. Holman, T. G. Nagaraja, Raghavendra Amachawadi, Samat Amat

## Abstract

*Trueperella pyogenes* is an important bacterial pathogen implicated in infections such as mastitis, metritis, pneumonia, and liver abscesses in both domestic and wild animals as well as endocarditis and prosthetic joint infections in humans. Understanding the genomic and metabolic features that enable *T. pyogenes* to colonize different anatomical sites within a host and its inter-kingdom transmission and survival is important for the effective control of this pathogen. We employed whole genome sequencing, phenotype microarrays, and antimicrobial susceptibility testing to identify genomic, metabolic and phenotypic features as well as antimicrobial resistance (AMR) genes in *T. pyogenes* recovered from different livestock, companion and wildlife animals. For comparative genomic analysis, 83 *T. pyogenes* genomes, including 60 isolated in the current study and 23 publicly available genomes were evaluated. These genomes represented *T. pyogenes* strains originated from 16 different body sites of 11 different animal hosts (e.g. bovine, swine, ovine, cervid, bison, equine, chamois, feline). Additionally, 49 *T. pyogenes* isolates (bovine, ovine, deer, swine and feline) were evaluated for phenotypic antimicrobial resistance using disk diffusion, and for metabolic profiling using the Biology GENIII MicroPlates. We identified that *T. pyogenes* strains are not host- or body site-specific. The presence of conserved virulence genes (*plo* and *fimA*), as well as genotypic and phenotypic AMR may contribute to *T. pyogenes’s* ability to cause infections in livestock, wildlife, and pets. Most of the tested isolates metabolized diverse carbon sources and chemical compounds, suggesting that this metabolic versatility may contribute to *T. pyogenes*’ survival, competitive advantage, and pathogenic potential.

**Importance:** *Trueperella pyogenes* is an important animal pathogen with zoonotic potential, posing a significant health concern to both animals and humans due to its ability to cause infections across different animal host species and tissues. Current understanding of this pathogen’s adaptability and survival mechanisms is limited. Here, we evaluated the genomic, virulence, metabolic, and antimicrobial resistance characteristics of *T. pyogenes* recovered from 16 different body sites of 11 different animal hosts (livestock, companion, and wild animals). We identified multiple antimicrobial resistance and virulence genes that may enable *T. pyogenes* for sustained infection and transmission. Additionally, *T. pyogenes* strains displayed metabolic versatility which could also contribute to its ability to thrive in diverse environments. Understanding the genomic and metabolic, and antimicrobial resistance characteristics that enable *T. pyogenes* to colonize different anatomical sites within a host and its transmission between different animal species is important for the effective control of this pathogen.

## Introduction

*Trueperella pyogenes* is a Gram-positive pleomorphic, non-motile, non-spore forming, and facultative anaerobic rod bacterium, formerly classified as *Arcanobacterium pyogenes* (Yassin et al., 2011). This bacterium is an important pathogen implicated in various infections in domestic and wild animals, including metritis, mastitis, and liver abscess in cattle, as well as pneumonia in swine (Ribeiro et al., 2015). Among livestock hosts, *T. pyogenes* is frequently isolated from cattle (both beef and dairy) and swine infections (Qin et al., 2022), but sheep, goats and horses can also be infected. In wild animals, *T. pyogenes*-associated chronic purulent infections (keratoconjunctivitis, brain and foot abscesses), pneumonia, abscesses, and necrobacillosis are common in captive and free-ranging white-tailed deer (N. W. Dyer et al., 2004; Rzewuska et al., 2019; K.-L. Zhao et al., 2011). *T. pyogenes* has also been isolated from the lungs of American bison (N. Dyer et al., 2013) big horn sheep (Garwood et al., 2020), and camels (Wareth et al., 2014), as well as the vaginal discharge of an okapi, and the kidneys of a royal python (Ahmed et al., 2020). *Trueperella pyogenes* is also a pathogenic for companion animals including cats and dogs (Wareth et al., 2018), with clinical manifestations ranging from cystitis and otitis externa, to pneumonia and wound infections (Billington et al., 2002; Hesselink & van den Tweel, 1990; Risseti et al., 2017). Recently, *T. pyogenes* has emerged as a zoonotic pathogen, causing clinical signs in humans including endocarditis, ulcers, and sepsis. The people affected by *T. pyogenes* were those working in close contact with farm animals and/or underlying conditions (Deliwala et al., 2020; Kavitha et al., 2010; Levy et al., 2009; Stuby et al., 2023). Thus, *T. pyogenes* is one of the important bacterial pathogens that can affect food-producing animals, companion animals, wildlife and humans.

The genomic and metabolic characteristics of *T. pyogenes* that enable colonization, spread, and infection of such a broad range of hosts and body sites are under-explored. Virulence factors could be one of the primary determinants of *T. pyogenes* multi-host and multi-niche colonization (K. Zhao et al., 2013). The main virulence factor possessed by this species is the cholesterol-dependent cytolysin exotoxin, called pyolysin, encoded by a gene, *plo*, which can damage host cell membranes through pore formation (Liang et al., 2022). Other virulence factors such as collagen-binding protein (*CbpA*), neuraminidases H and P (*NanH* and *NanP*), and types A, C, E, and G, fimbriae, encoded by *FimA, FimC, FimE*, and *FimG* genes, can contribute to pathogenesis by enabling colonization of different mucosal niches (Billington et al., 1997; Jost et al., 2001; Pietrocola et al., 2007; Rzewuska et al., 2019). Niche-specific genetic, metabolic and phenotypic evolution has been relatively well documented in some bacterial species, particularly *Escherichia coli* (Barua et al., n.d.; Favate et al., 2023; Hernández-Chiñas et al., 2023; Yu et al., 2021), and *Lactobacillus* spp. (Baek et al., 2023; Pan et al., 2021; Zhang et al., 2020). Thus, it is likely that *T. pyogenes* strains in different mammalian hosts and different body sites within an animal may have evolved both genetically and phenotypically to adapt to these different niches. However, there is limited data available on the genomic (diverse pangenome and intergenotypic variability), phenotypic (metabolic pathways), and ecotypic (host-/niche-specificity) diversity among *T. pyogenes* present in different hosts, and body sites.

Resistance to antimicrobials commonly used in animal production systems, such as tetracyclines (Yapicier et al., 2022), trimethoprim-sulfamethoxazole, and macrolides-lincosamides-streptogramin B (MLS_B_) has been reported in *T. pyogenes* isolates (Kwiecień et al., 2021). However, whether the antimicrobial susceptibility of *T. pyogenes* strains varies depending on the host and niche-origin is unknown. Given that *T. pyogenes* has the potential to transfer among domestic (including livestock and pets) and wild animals, and from animals to humans, and it is important from a One Health perspective to characterize antimicrobial resistance (AMR) in *T. pyogenes* originated from different hosts. Thus, the objectives of this study were to: 1) isolate *T. pyogenes* from different animal species (domestic and wild) and infection body sites, 2) perform comparative genomic and metabolic analyses to identify genotypic and phenotypic characteristics associated with host-/niche specificity, and 3) characterize and compare genotypic and phenotypic AMR profiles among *T. pyogenes* isolated from different animal hosts and body sites.

## Results

### *T. pyogenes* strains origin

The 60 *T. pyogenes* strains used in the present study were isolated from bovine (n = 49), swine (n =3), ovine (n =3), cervid (n = 3), bison (n = 1), feline (n = 1) animal host species, with strains originating most frequently from ruminal tissues (n = 24) and lungs (n = 22). Of note, one of the three swine-orign isolates was obtained from ATCC (strain# ATCC-19411). Additionally, 23 publicly available genomes from bovine (n = 9), swine (n = 8), buffalo (n = 2), canine (n = 1), caprine (n = 1), chamois (n = 1), and equine (n = 1) were also included (Table 1).

**Table 1.** *T. pyogenes* genome assemblies and isolates.

| Accession number | Isolate ID | Animal host | Anatomical body site | Source | AST | Metabolite |
| --- | --- | --- | --- | --- | --- | --- |
| GCF_000599565 | MS249 | Bovine | Uterus | NCBI | No | No |
| GCF_000612055 | TP6375 | Bovine | Uterus | NCBI | No | No |
| GCF_001281085 | 2012CQ-ZSH | Caprine | Lung | NCBI | No | No |
| GCF_002071825 | UFV1 | Bovine | Uterus | NCBI | No | No |
| GCF_002762575 | Bu5 | Buffalo | Wound | NCBI | No | No |
| GCF_003055835 | Arash114 | Buffalo | Uterus | NCBI | No | No |
| GCF_003076295 | TP4479 | Swine | N/A | NCBI | No | No |
| GCF_003076315 | TP-2849 | Swine | Lung | NCBI | No | No |
| GCF_003971445 | TP1 | Bovine | Lung | NCBI | No | No |
| GCF_003971465 | TP2 | Bovine | Joint | NCBI | No | No |
| GCF_003971485 | TP3 | Swine | Lung | NCBI | No | No |
| GCF_003971545 | TP4 | Swine | Lung | NCBI | No | No |
| GCF_004123715 | SH03 | Swine | Lung | NCBI | No | No |
| GCF_004009995.2 | SH02 | Swine | Lung | NCBI | No | No |
| GCF_004564055 | SH01 | Swine | Lung | NCBI | No | No |
| GCF_012109075 | jx18 | Swine | Lung | NCBI | No | No |
| GCF_024741855 | 22/KM0800 | Bovine | Wound | NCBI | No | No |
| GCF_024803545 | EMSSI21 | Bovine | Wound | NCBI | No | No |
| GCF_024803565 | EMSSI54 | Bovine | Wound | NCBI | No | No |
| GCF_024803585 | EMSSI48 | Bovine | Wound | NCBI | No | No |
| GCF_030078095 | 09KM1269 | Canine | N/A | NCBI | No | No |
| GCF_030078115 | 13KM1326 | Equine | N/A | NCBI | No | No |
| GCF_030078145 | 13OD0707 | Chamois | N/A | NCBI | No | No |
| JBAGLQ000000000 | ATCC_19411 | Swine | N/A | This study | No | Yes |
| JBAGLR000000000 | 93CBB | Bovine | Vaginal swab | This study | Yes | Yes |
| JBAGLS000000000 | 51CBC | Bovine | Vaginal swab | This study | Yes | Yes |
| JBAGLT000000000 | 4618_1 | Bovine | Abscess | This study | Yes | Yes |
| JBAGLU000000000 | 23_564_0001_02 | Bovine | Liver | This study | No | Yes |
| JBAGLV000000000 | 23_50711_2 | Bovine | Lung | This study | No | Yes |
| JBAGLW000000000 | 23_4130 | Bovine | Lung | This study | Yes | Yes |
| JBAGLX000000000 | 23_30761_2 | White-tailed deer | Lung | This study | Yes | Yes |
| JBAGLY000000000 | 22_7871 | Ovine | Mammary | This study | Yes | Yes |
| JBAGLZ000000000 | 22_5896 | Bovine | Lung | This study | Yes | Yes |
| JBAGMA000000000 | 22_5293 | Bovine | Foot | This study | Yes | Yes |
| JBAGMB000000000 | 22_5283 | Bovine | Lung | This study | Yes | Yes |
| JBAGMC000000000 | 22_4838 | Bovine | Lung | This study | Yes | Yes |
| JBAGMD000000000 | 22_4813 | Bovine | Lung | This study | No | Yes |
| JBAGME000000000 | 22_2570 | Bovine | Milk | This study | No | Yes |
| JBAGMF000000000 | 22_2565 | Bison | Lung | This study | No | No |
| JBAGMG000000000 | 22_2564 | Bovine | Peri fluid | This study | No | Yes |
| JBAGMH000000000 | 22_11656 | Bovine | Lung | This study | Yes | Yes |
| JBAGMI000000000 | 22_11177 | Bovine | Lung | This study | Yes | Yes |
| JBAGMJ000000000 | 22_11067 | Ovine | Umbilicus swab | This study | Yes | Yes |
| JBAGMK000000000 | 22_10600 | Feline | Nasal swab | This study | Yes | Yes |
| JBAGML000000000 | 21_2041 | White-tailed deer | Lung | This study | Yes | Yes |
| JBAGMM000000000 | 2023_4_80 | Bovine | Rumen tissue | This study | Yes | Yes |
| JBAGMN000000000 | 2023_3_72 | Bovine | Rumen tissue | This study | Yes | Yes |
| JBAGMO000000000 | 2023_3_71 | Bovine | Rumen tissue | This study | No | No |
| JBAGMP000000000 | 2023_3_67 | Bovine | Rumen tissue | This study | No | Yes |
| JBAGMQ000000000 | 2023_3_60 | Bovine | Rumen tissue | This study | No | No |
| JBAGMR000000000 | 2023_3_4 | Bovine | Rumen tissue | This study | Yes | Yes |
| JBAGMS000000000 | 2023_3_235 | Bovine | Rumen tissue | This study | Yes | Yes |
| JBAGMT000000000 | 2023_3_175 | Bovine | Rumen tissue | This study | No | Yes |
| JBAGMU000000000 | 2023_3_174 | Bovine | Rumen tissue | This study | No | No |
| JBAGMV000000000 | 2023_3_167 | Bovine | Rumen tissue | This study | Yes | No |
| JBAGMW000000000 | 2023_3_164 | Bovine | Rumen tissue | This study | No | Yes |
| JBAGMX000000000 | 2023_3_163 | Bovine | Rumen tissue | This study | No | Yes |
| JBAGMY000000000 | 2023_3_119 | Bovine | Rumen tissue | This study | No | No |
| JBAGMZ000000000 | 2023_3_111 | Bovine | Rumen tissue | This study | No | No |
| JBAGNA000000000 | 2022_1_96 | Bovine | Rumen tissue | This study | No | No |
| JBAGNB000000000 | 2022_1_94 | Bovine | Rumen tissue | This study | No | Yes |
| JBAGNC000000000 | 2022_1_79 | Bovine | Rumen tissue | This study | No | Yes |
| JBAGND000000000 | 2022_1_77 | Bovine | Rumen tissue | This study | Yes | No |
| JBAGNE000000000 | 2022_1_75 | Bovine | Rumen tissue | This study | Yes | No |
| JBAGNF000000000 | 2022_1_30 | Bovine | Rumen tissue | This study | Yes | Yes |
| JBAGNG000000000 | 2022_1_29 | Bovine | Rumen tissue | This study | Yes | Yes |
| JBAGNH000000000 | 2022_1_27 | Bovine | Rumen tissue | This study | No | Yes |
| JBAGNI000000000 | 2022_1_14 | Bovine | Rumen tissue | This study | No | Yes |
| JBAGNJ000000000 | 8896 | Swine | Lung | This study | Yes | No |
| JBAGNK000000000 | 7762 | Bovine | Lung | This study | Yes | No |
| JBAGNL000000000 | 1510 | Bovine | Lung | This study | Yes | No |
| JBAGNM000000000 | 1494 | White-tailed deer | Lung | This study | Yes | No |
| JBAGNN000000000 | 1446 | Bovine | Lung | This study | Yes | No |
| JBAGNO000000000 | 1409 | Bovine | Lung | This study | Yes | No |
| JBAGNP000000000 | 1361 | Bovine | Lung | This study | Yes | Yes |
| JBAGNQ000000000 | 1355 | Ovine | Lung | This study | Yes | No |
| JBAGNR000000000 | 1259 | Bovine | Lung | This study | Yes | No |
| JBAGNS000000000 | 315 | Bovine | Liver abscess | This study | Yes | No |
| JBAGNT000000000 | 311 | Bovine | Liver abscess | This study | Yes | No |
| JBAGNU000000000 | 306 | Bovine | Liver abscess | This study | Yes | No |
| JBAWKN000000000 | 23_2752 | Swine | Lung | This study | Yes | Yes |
| JBAWKO000000000 | 2023_3_104 | Bovine | Rumen tissue | This study | No | Yes |
| JBAWKP000000000 | 5861 | Bovine | Lung | This study | Yes | No |

### Genome assembly and annotation

Of the 60 *T. pyogenes* genomes that we sequenced (59 new strains and 1 from ATCC), the average genome size was 2,278,603 ± 6,648 (SE) bp, G+C content was 59.6 ± 0.01%, and the number of coding sequences was 2,026 ± 6.51 (Table S1). These genomes contained 46 ± 0.05 tRNA genes, 3 rRNA genes, and 814 ± 8.6 unclassified hypothetical protein coding regions on average. A total of 54 strains (90.0%) had the CRISPR-associated nuclease/helicase gene *cas3*, and 56 strains (93.3%) displayed at least one repeat region in their genomes (Table S1).

### Comparative genome analysis

In addition to the genomes of the 60 strains that we sequenced, 23 publicly available genome sequences were included in our comparative genomic analysis to gain more comprehensive insight into the *T. pyogenes* genomes. The pangenome of 83 *T. pyogenes* genomes contained 5,612 genes, and of these, 1,470 were core genes that were shared among all genomes. The remaining 4,142 genes were accessory genes that can be divided into soft core (95% to 99% of genomes; n = 155), shell (15% to 95% of genomes; n = 609), and cloud (0% to 15% of genomes; n = 3,378) genes. *T. pyogenes* appears to have an open pangenome as there was no plateau in the number of new genes added to the pangenome as more genomes were included (Fig. 1). Phylogenetic analysis showed that there were several clades with bovine and swine isolates largely clustering separately from each other (Fig. 2). Moreover, there was no distinct clustering of genomes based on the anatomical body site or country of origin. There were 20 genomes with an average nucleotide identity (ANI) of > 99.99% with at least one other genome, and 22 with an ANI between 99.5-99.99% (Table S2). Based on the phylogenetic tree and their ANI values, there were certain *T. pyogenes* isolates recovered from different host species that appeared to be nearly identical (ANI > 99.99%). This included the isolates 8896 and 1510, which were recovered from the lungs of a pig and cattle in 2016 and 2023, respectively, at the NDSU-VDL. *T. pyogenes* 4618_1 (bovine abscess) and 23_2752 (swine lung) were also isolated in 2016 and 2023, respectively, at the NDSU-VDL, suggesting possible inter-species transfer (Table S2). Consequently, it appears that these strains can colonize and cause disease in both host species, in addition to persisting in the animal population/environment for long periods of time. Additionally, some isolates could be considered more potentially pathogenic or virulent, such as isolates 2022_1_29 and 2022_1_77 from bovine rumen tissue. These isolates share eight ARGs, resulting in genotypical AMR to six antimicrobial classes (i.e. aminoglycosides, biocides, glycopeptides, MLSb, sulfonamides, and tetracyclines) and phenotypical resistance to clindamycin (lincosamide), gentamicin (aminoglycoside), and tetracycline (Table S4). They are also genetically similar, with ANI > 99.99% (Table S2), suggesting that they are form the same strain of *T. pyogenes*.

**Fig. 1.**
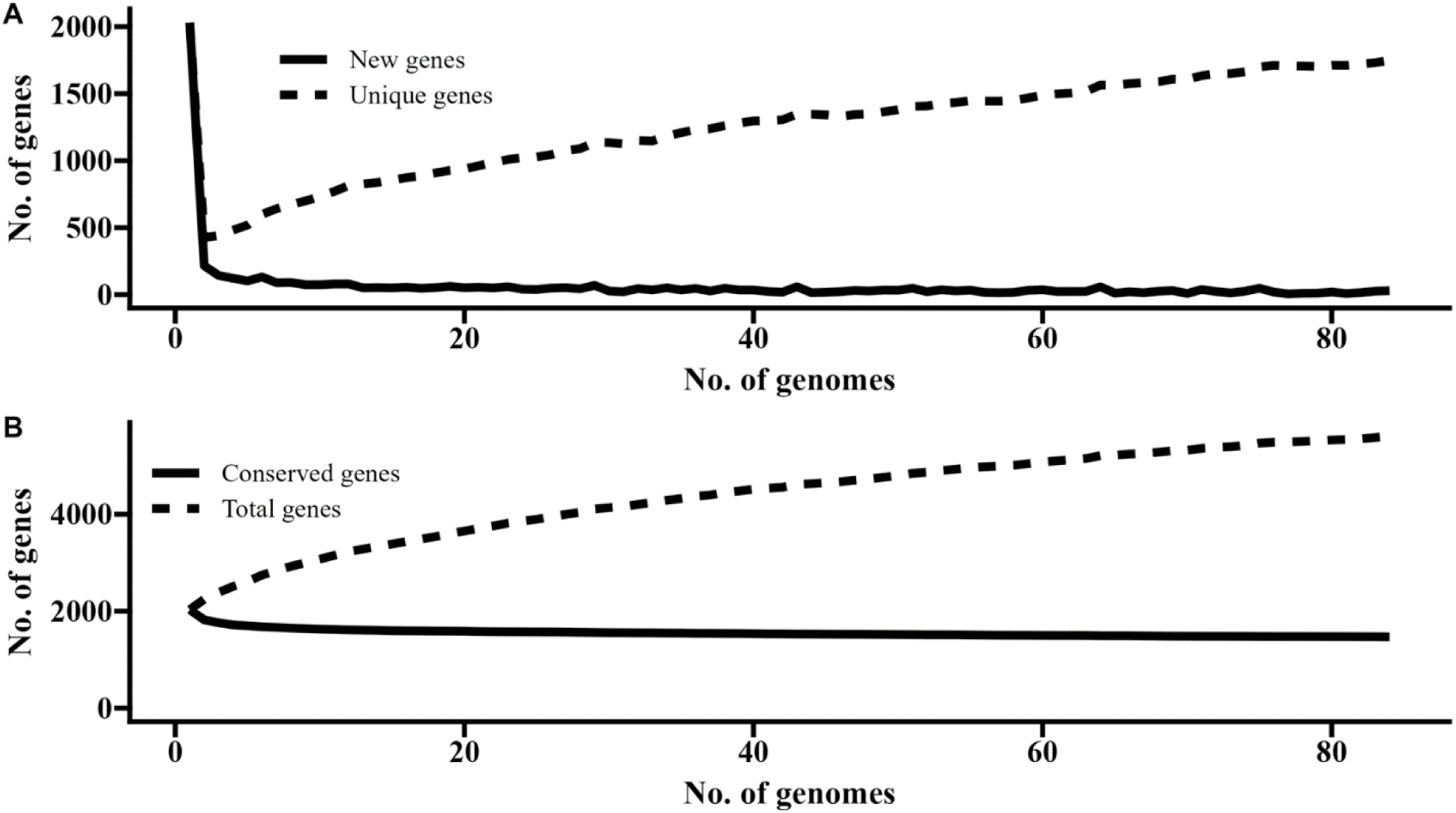
Distribution of A) unique and B) conserved genes in the pangenome of *T. pyogenes* (n = 83)

**Fig 2.**
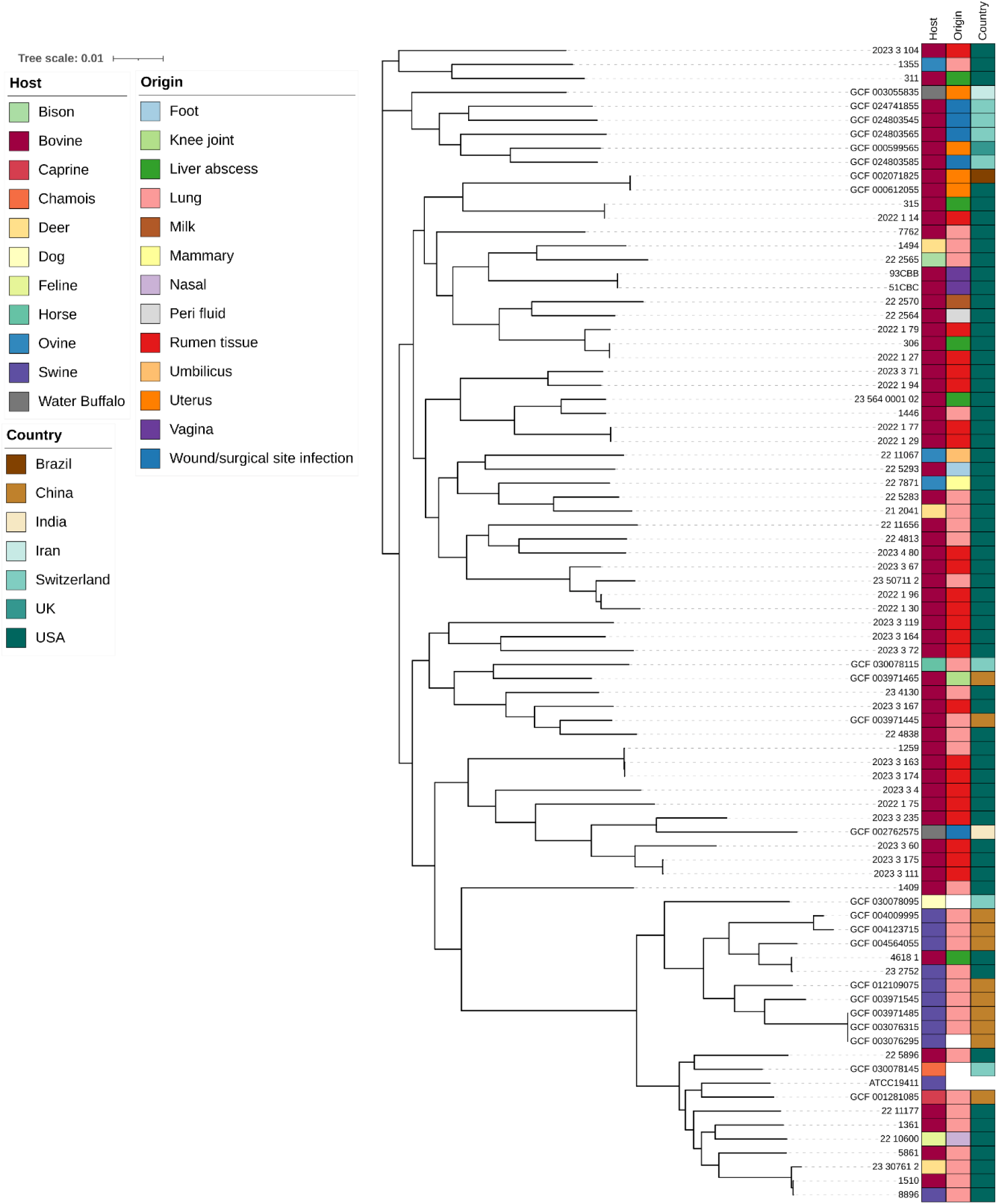
Maximum likelihood phylogenetic tree of *Trueperella pyogenes* isolates obtained from this study (n = 60) and publicly available genomes (n = 23). Phylogeny was inferred from the alignment of 1,470 core genes. The scale bar represents substitutions per nucleotide.

### Virulence genes and genotypes

The 83 *T. pyogenes* genomes investigated in this study were screened for the presence of 7 known virulence genes. The most prevalent virulence genes were *plo* (98.8%), *fimA* (97.6%), *fimE* (90.4%), and *nanP* (63.9%), followed by *fimC* (51.8%), *cbpA* (41%), and *nanH* (14.5%) (Fig. 3). Based on the combination of virulence genes, the genomes were categorized into 20 genotypes. The predominant genotypes were VIII with the *plo, fimA, fimC, fimE,* and *nanP*, genes (14.5%), XVI with the *plo*, *fimA*, *fimE, nanP* genes (14.5%), XI with the *plo*, *fimA*, *fimE, nanP*, and *cbpA* genes (9.6%), VI with the *plo*, *fimA*, *fimC, fimE*, and *cbpA* genes (7.2%), and XIII with *plo, fimA, fimC*, and *fimE* (7.2%) (Table S3). The other genotypes were less abundant, and present in less than 7% of the genomes each. The proportion of virulence genes among different anatomical body sites or between animal host species did not follow any detectable patterns and it is displayed on Fig. 3.

**Fig. 3.**
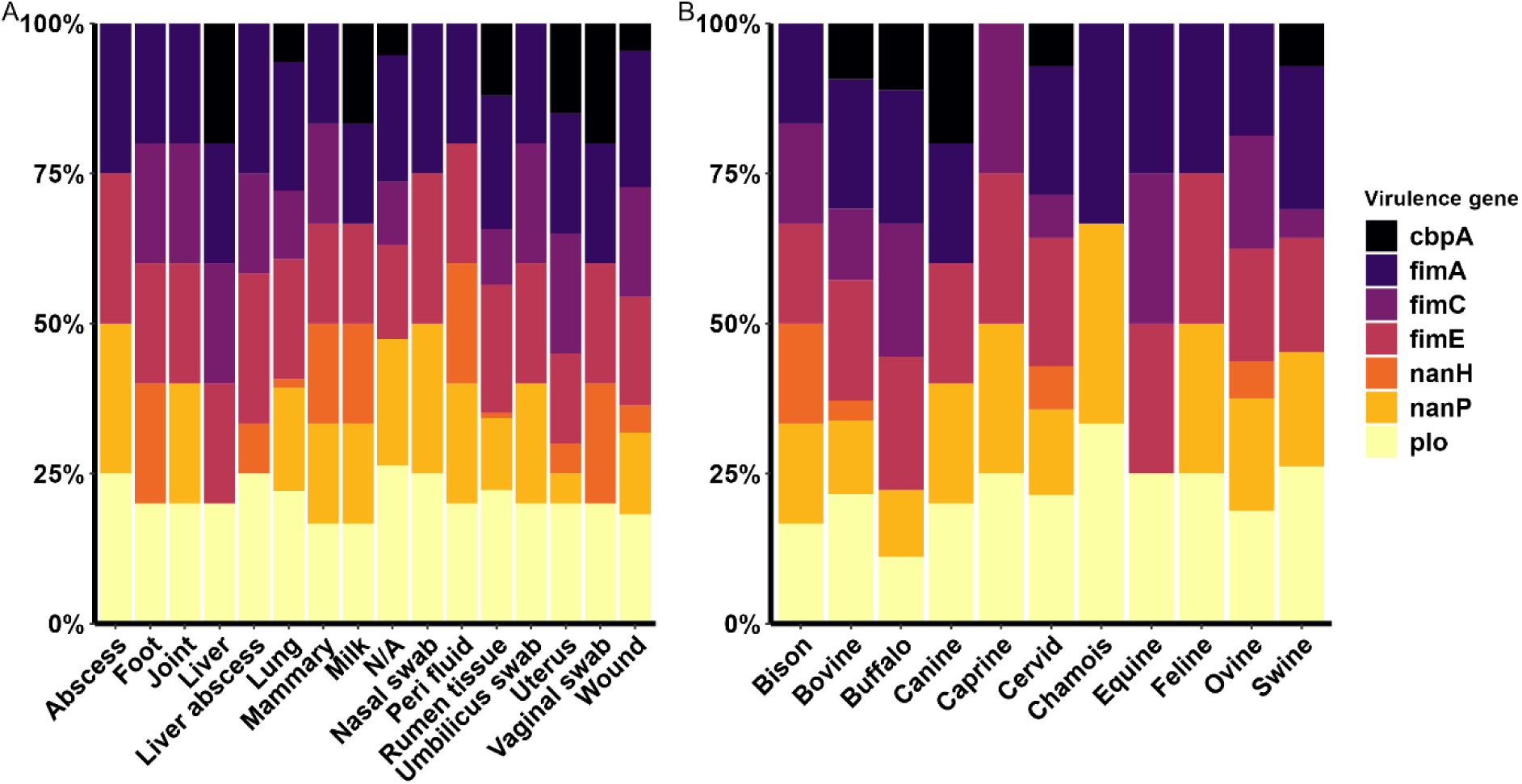
Distribution and proportion of virulence genes in the genomes of *T. pyogenes* isolates by A) infection site and B) host species.

### Antimicrobial resistance gene profiles

There were 19 different antimicrobial resistance genes (ARGs) identified among the 83 *T. pyogenes* genomes (Table 2). The ribosomal protection protein gene *tet(*W/32/O) conferring resistance to the tetracyclines was the most prevalent ARG (53% of genomes), followed by *erm*(X) (31.3%), *vanG* (25.3%), *sul1* (21.7%), and *qacEdelta1* (20.5%) (Table 2). Moreover, 19 *T. pyogenes* genomes (22.9%) did not carry any ARGs, including the ATCC 19411 reference strain. The ARGs identified confer resistance to many different antimicrobials classes, namely tetracycline (69.9%), MLS_B_ (31.3%), glycopeptides (25.3%), sulfonamides (21.7%), biocides (20.5%), aminoglycosides (18.1%), and phenicols (3.6%) (Table 2). When the *T. pyogenes* isolates were grouped based on body site origin, the liver/liver abscess isolates had a similar ARGs profile with genes encoding resistance to tetracyclines, aminoglycosides, and MLS_B_. This is similar to the ARGs observed in the genomes of multiple ruminal tissue associated isolates (Fig. 4). Several isolates encoded multiple ARGs that appeared to be on the same contig together with certain mobile genetic element genes (Fig. S1). *T*. *pyogenes* 2023_3_67, isolated from bovine ruminal tissue, carried eight ARGs, including *erm*(X) and *tet*(33) genes were co-located on a 8,048 bp contig with a *repA* gene (replication initiation) and *ant(3’’)-IIa*, *qacEdelta1*, *sul1* on a 4.214 bp contig. In two other bovine ruminal tissue isolates (2022_1_29 and 2022_1_75), *sul1*, *qacEdelta1*, *aadA2*, and *ant(2’’)-Ia* were found together on a 5,857 bp contig also containing the plasmid-associated genes *parA* (partition) and *int* (integrase). The *sul1* and *qacEdelta1* genes were linked together in all isolates carrying both ARGs. Additionally, isolates 2022_1_29 and 2022_1_77, also recovered from bovine rumen tissue, shared eight ARGs conferring resistance to six antimicrobial classes (i.e. aminoglycosides, biocides, glycopeptides, MLSB, sulfonamides, and tetracyclines). They were also genetically very similar (ANI > 99.99%; Table S2), suggesting that they are the same strain.

**Fig. 4.**
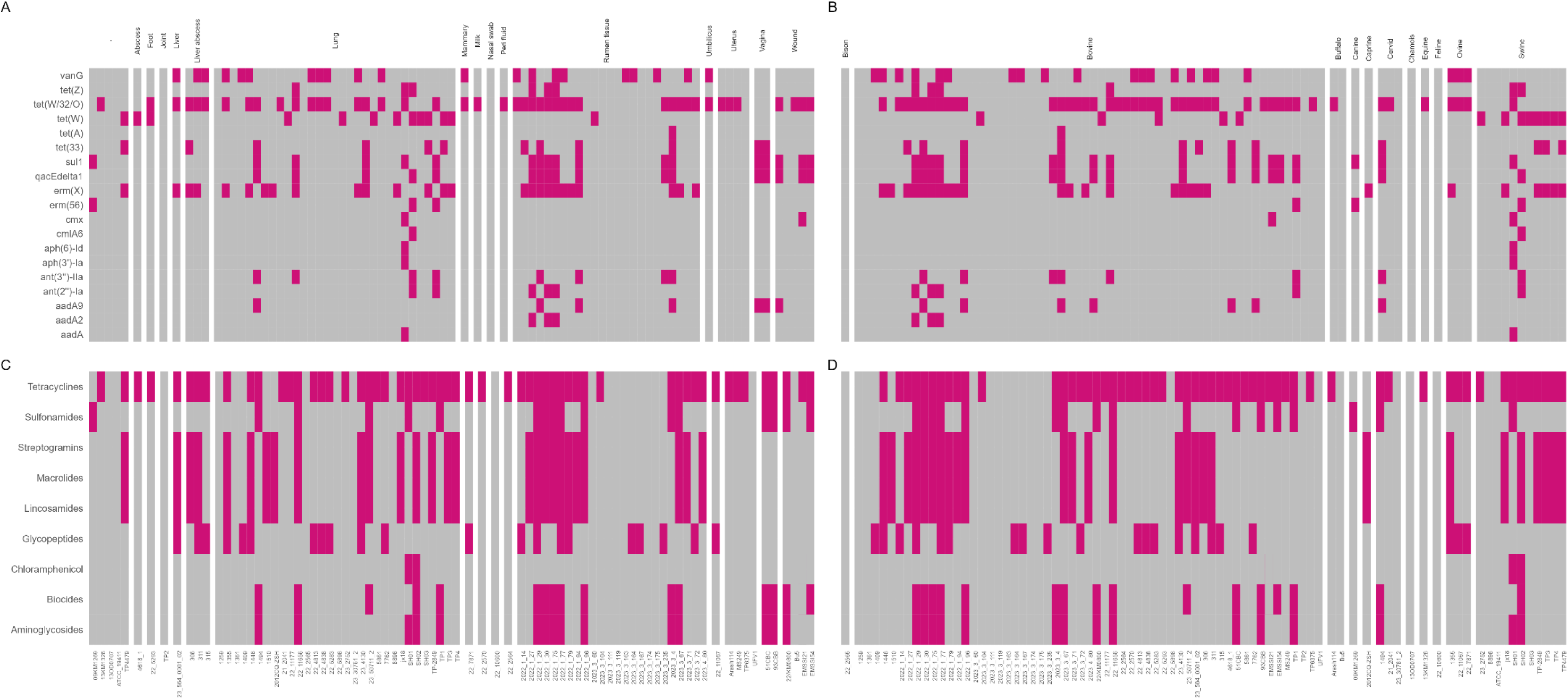
Antimicrobial resistance genes (ARGs) in *T. pyogenes* isolates (n = 83) by A) body site, and B) host species, and antimicrobial class for the ARGs confer resistance to by C) anatomical body site, and D) host species.

**Table 2.**
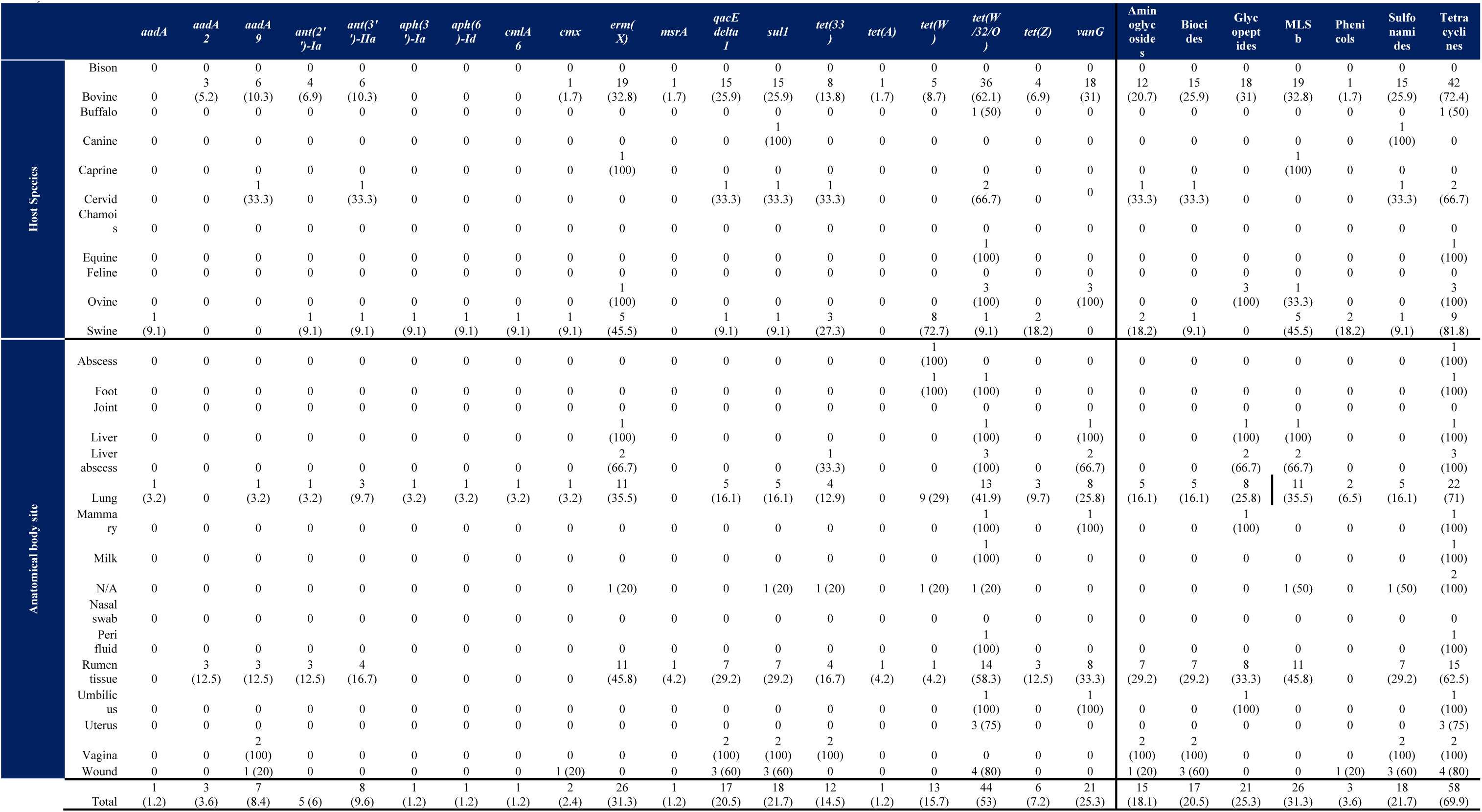
Distribution (%) of antimicrobial resistance genes in the genomes of *T. pyogenes* from different host species and anatomical body sites (n = 83)

### Phenotypic antimicrobial susceptibility

The zone of inhibition (ZOI; in mm) produced by each of the antibiotic disks tested ranged from 0-37 (21.1 ± 2.3 [SEM]) for clindamycin, 0-53 (32.2 ± 2.8) for erythromycin, 15-41 (35.8 ± 0.6) for chloramphenicol, 12-23 (16.4 ± 0.3) for ciprofloxacin, 0-30 (23.4 ± 0.9) for gentamicin, 38-60 (52.3 ± 0.7) for penicillin, 31-43 (38.0 ± 0.4) for sulfamethoxazole/trimethoprim, 9-46 (20.1 ± 1.7) for tetracycline, and 28-34 (21.1 ± 0.2) for vancomycin (Table S4). The ZOI was largest for penicillin, followed by erythromycin, sulfamethoxazole/trimethoprim, chloramphenicol, and vancomycin. Certain isolates (i.e. isolates 2022_1_96 and 2022_1_29 from bovine rumen tissue, and 23_50711 from bovine lung) were also completely resistant (0 mm ZOI) to gentamicin, clindamycin, or erythromycin (Fig. 5).

**Fig. 5.**
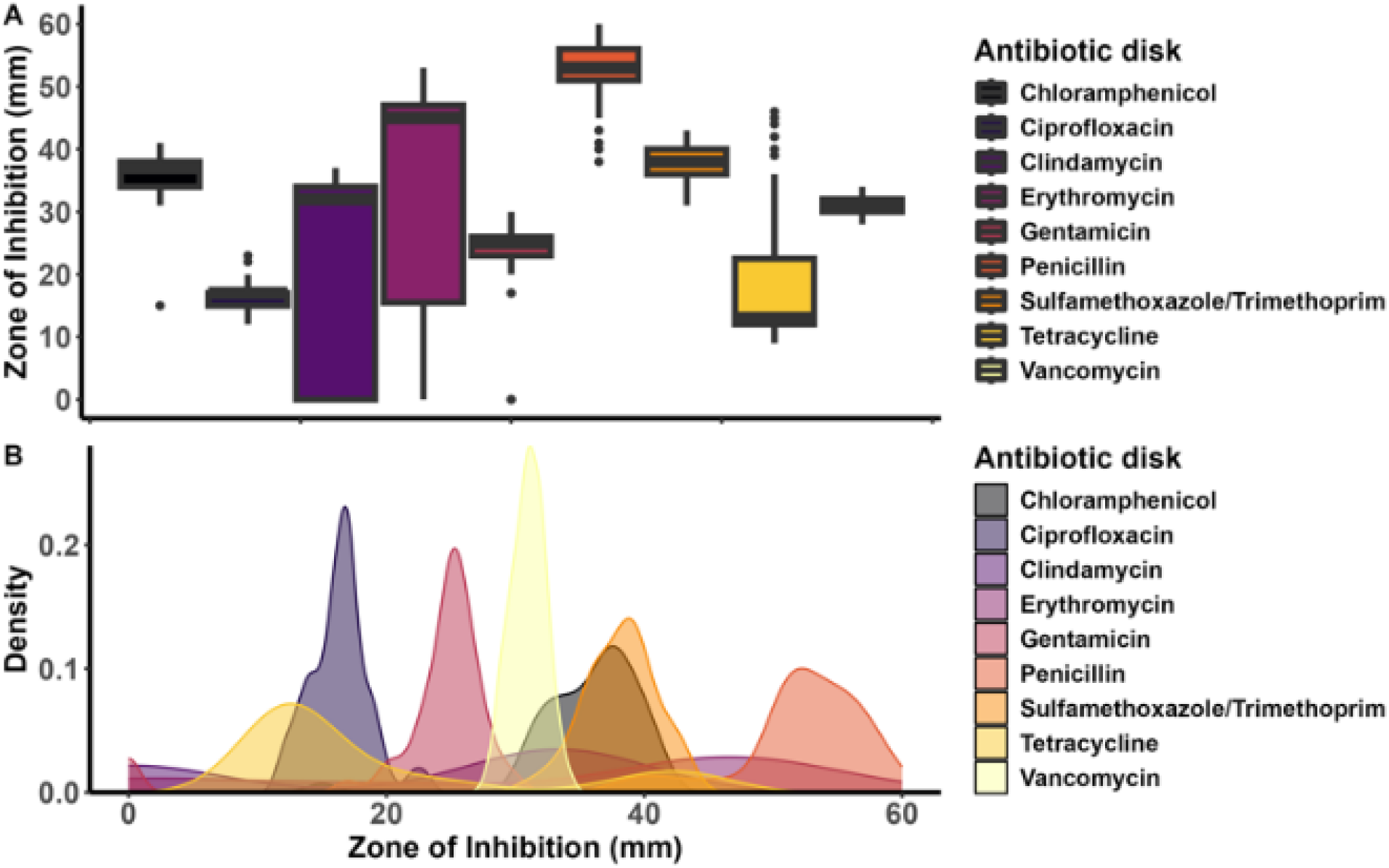
Differences in the A) variation and B) density of ZOI diameters (mm) for each tested antibiotic across selected *T. pyogenes* isolates (n = 49)

According to the breakpoints available for *Streptococcus pneumoniae,* the majority of *T. pyogenes* isolates were resistant to tetracycline (75%), followed by clindamycin (37.5%), erythromycin (24.5%), gentamicin (6.1%), and chloramphenicol (2.1%) (Table 3). None of the *T. pyogenes* isolates tested were resistant to ciprofloxacin, sulfamethoxazole/trimethoprim, penicillin, or vancomycin. Concomitantly, 49% of isolates had intermediate resistance to ciprofloxacin, 6.1% to erythromycin, and 4.2% to tetracycline (Table 3). When comparing different animal hosts and body sites, bovine isolates were more frequently resistant to tetracycline, erythromycin, clindamycin, ciprofloxacin, and chloramphenicol than the other animal hosts combined. Additionally, *T. pyogenes* isolates from the ruminal tissue were mostly resistant to erythromycin, clindamycin, ciprofloxacin, and chloramphenicol than the lung and all the other body sites-associated isolates (Fig. 6).

**Fig. 6.**
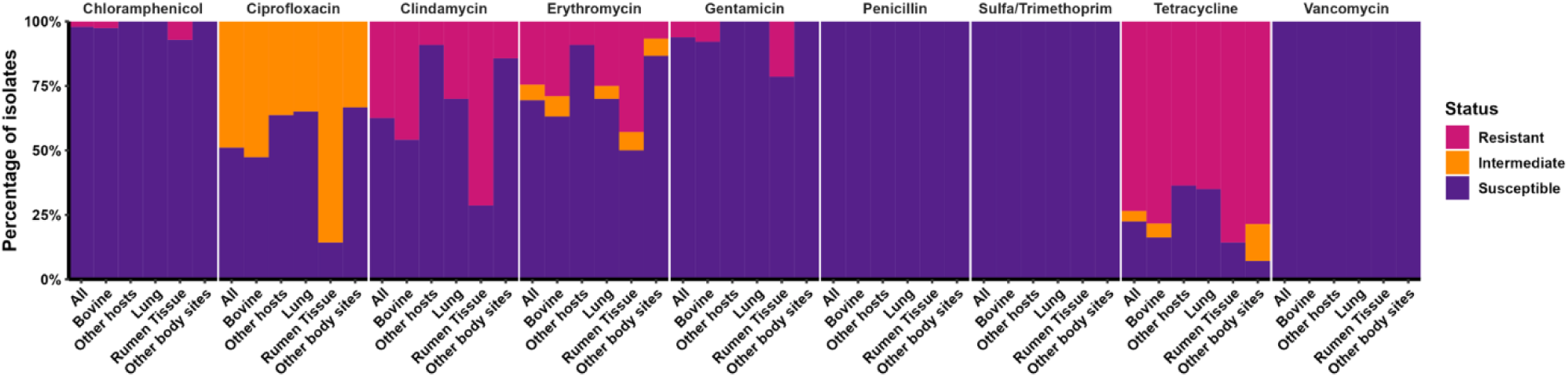
The proportion of resistant, intermediate, and susceptible *T. pyogenes* isolates originating from different animal hosts and body sites based on disk diffusion (n = 49)

**Table 3.** Disk diffusion zone of inhibition (ZOI) and phenotypical antibiotic resistance *T*. *pyogenes* isolates against selected antibiotics (n = 49)

| Antibiotic disk | Zone diameter (mm) |  |  |  |  |  |  |  |  | Phenotype |  |  |
| --- | --- | --- | --- | --- | --- | --- | --- | --- | --- | --- | --- | --- |
|  | 0-5 | 6-10 | 11-15 | 16-20 | 21-25 | 26-30 | 31-35 | 36-40 | >40 | S | I | R |
| Gentamicin<sup>3</sup> | 6.1% | - | - | 6.1% | 51.0% | 36.7% | - | - | - | 46 (93.9%) | 0 | 3 (6.1%) |
| Ciprofloxacin<sup>1</sup> | - | - | 30.6% | 65.3% | 4.1% | - | - | - | - | 25 (51%) | 24 (49%) | 0 |
| Erythromycin<sup>2</sup> | 20.4% | 2.0% | 2.0% | 6.1% | 8.2% | - | - | - | 61.2% | 34 (69.4%) | 3 (6.1%) | 12 (24.5%) |
| Sulfamethoxazole /Trimethoprim<sup>2</sup> | - | - | - | - | - | - | 16.3% | 67.3% | 16.3% | 49 (100%) | 0 | 0 |
| Clindamycin<sup>2</sup> | 35.4% | 2.1% | - | - | - | 4.2% | 47.9% | 10.4% | - | 30 (62.5%) | 0 | 18 (37.5%) |
| Chloramphenicol<sup>2</sup> | - | - | 2.1% | - | - | - | 35.4% | 56.3% | 6.3% | 47 (97.9%) | 0 | 1 (2.1%) |
| Penicillin<sup>2</sup> | - | - | - | - | - | - | - | 4.1% | 95.9% | 49 (100%) | 0 | 0 |
| Vancomycin<sup>2</sup> | - | - | - | - | - | 34.7% | 65.3% | - | - | 49 (100%) | 0 | 0 |
| Tetracycline<sup>2</sup> | - | 6.3% | 52.1% | 12.5% | 6.3% | 2.1% | - | 20.8% | - | 10 (20.8%) | 2 (4.2%) | 36 (75%) |

Vertical red lines represent the breakpoints for the tested antibiotic disks. <sup>1</sup>Coryneform breakpoints. <sup>2</sup>*Streptococcus pneumoniae* breakpoints. <sup>3</sup>ZOI = 0 resistance breakpoint.
S, susceptible; I, intermediate; R, resistant

### Comparison between genotypic and phenotypic antimicrobial susceptibility

We sought to identify whether and to what extent the presence of an ARG influenced phenotypic AMR in *T. pyogenes*. For this, the associations between ARGs and the ZOI (mm) obtained from the disk diffusion assay were determined with the Kruskal-Wallis test. The presence of an ARG in the isolate was significantly associated with the size of the ZOI (Fig. 7). Isolates that carried ARGs conferring resistance to clindamycin, erythromycin, gentamicin, sulfamethoxazole/trimethoprim, or tetracycline had a significantly (*P* < 0.05) smaller ZOI than those isolates that lacked ARGs. The ZOI for vancomycin was similar (*P* = 0.49) between isolates that had *vanG* versus the ones that didn’t (Fig. 7), again demonstrating that *vanG* provides only low-level resistance to vancomycin. Additionally, the correlation between genotypical and phenotypical resistance was assessed using either the chi-square or Fisher’s exact tests. There was a positive correlation (*P* < 0.05) between genotypic and phenotypic resistance for aminoglycosides, macrolides, tetracyclines, sulfonamides, and lincosamides (Fig. 7), meaning that if an isolate was predicted to be resistant based on its genotype, then it was likely phenotypically resistant as well. Generally, *T. pyogenes* isolates are less susceptible to tetracycline and macrolides, while they are mostly susceptible to penicillin, vancomycin, sulfamethoxazole/trimethoprim, and gentamycin.

**Fig. 7.**
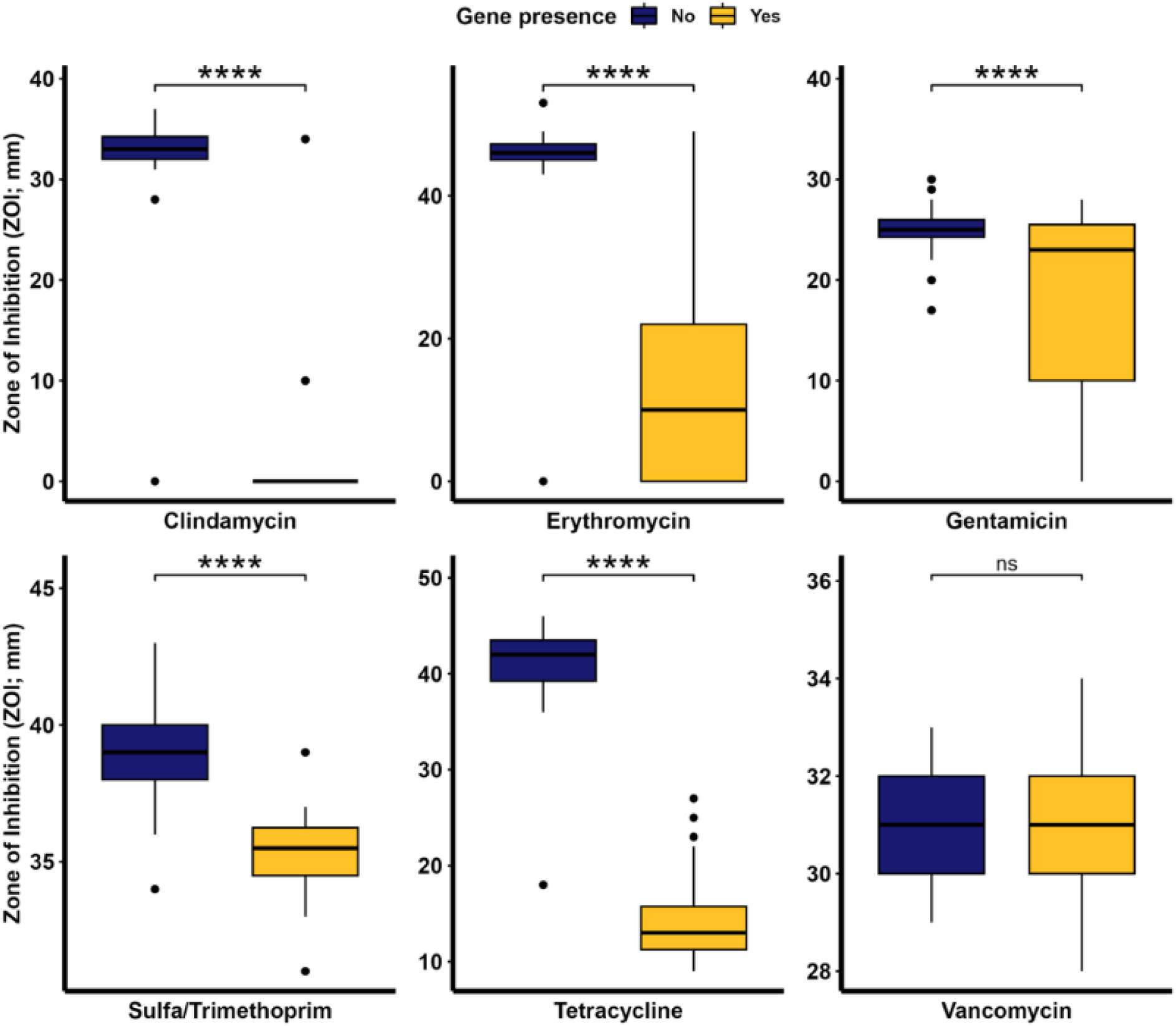
Disk diffusion zone of inhibition (mm) for antimicrobials against *Trueperella pyogenes* isolates (n = 49) based on the presence or absence of corresponding antimicrobial resistance genes in their genomes. **** indicates a significant difference with *P* -value < 0.001 and ns indicates no significance (*P* -value > 0.05). Of note, the diameter of the disk was 6 mm.

### Metabolic characterization of selected *T. pyogenes*

To identify whether there is a common metabolic signature among *T. pyogenes* strains originating from different body sites within an animal, and from different animal hosts, we used Biology GENIII MicroPlates to biochemically profile 49 *T. pyogenes* isolates. Each strain was tested in 96-well Microplate containing 71 carbon sources and 23 chemical sensitivity analysis (Table S5). After 48 h of incubation, all *T. pyogenes* isolates tested turned purple in the positive control, and no color changed in the negative control well based on the OD_630_. Most isolates grew well at pH = 6.0 and low salt concentration (up to 4% NaCl) but not in under acidic conditions (pH = 5) and higher salt concentrations (8% NaCl). The carbon sources readily utilized by all or most of the isolates were α-D-lactose, N-acetyl neuraminic acid, and α-D-glucose (Fig. 8). Conversely, some carbon sources were not preferred by most isolates, but were metabolized by 1 to 3 isolates after a longer lag phase (Fig. 9). Most or all isolates grew normally and were able to metabolize 1% sodium lactate, tetrazolium violet, tetrazolium blue, nalidixic acid, lithium chloride, potassium telluride, aztreonam, and sodium butyrate. No specific differences between isolates of different animal hosts or body sites on their growth with various carbon sources or chemicals was observed in this biochemical microplate panel.

**Fig. 8.**
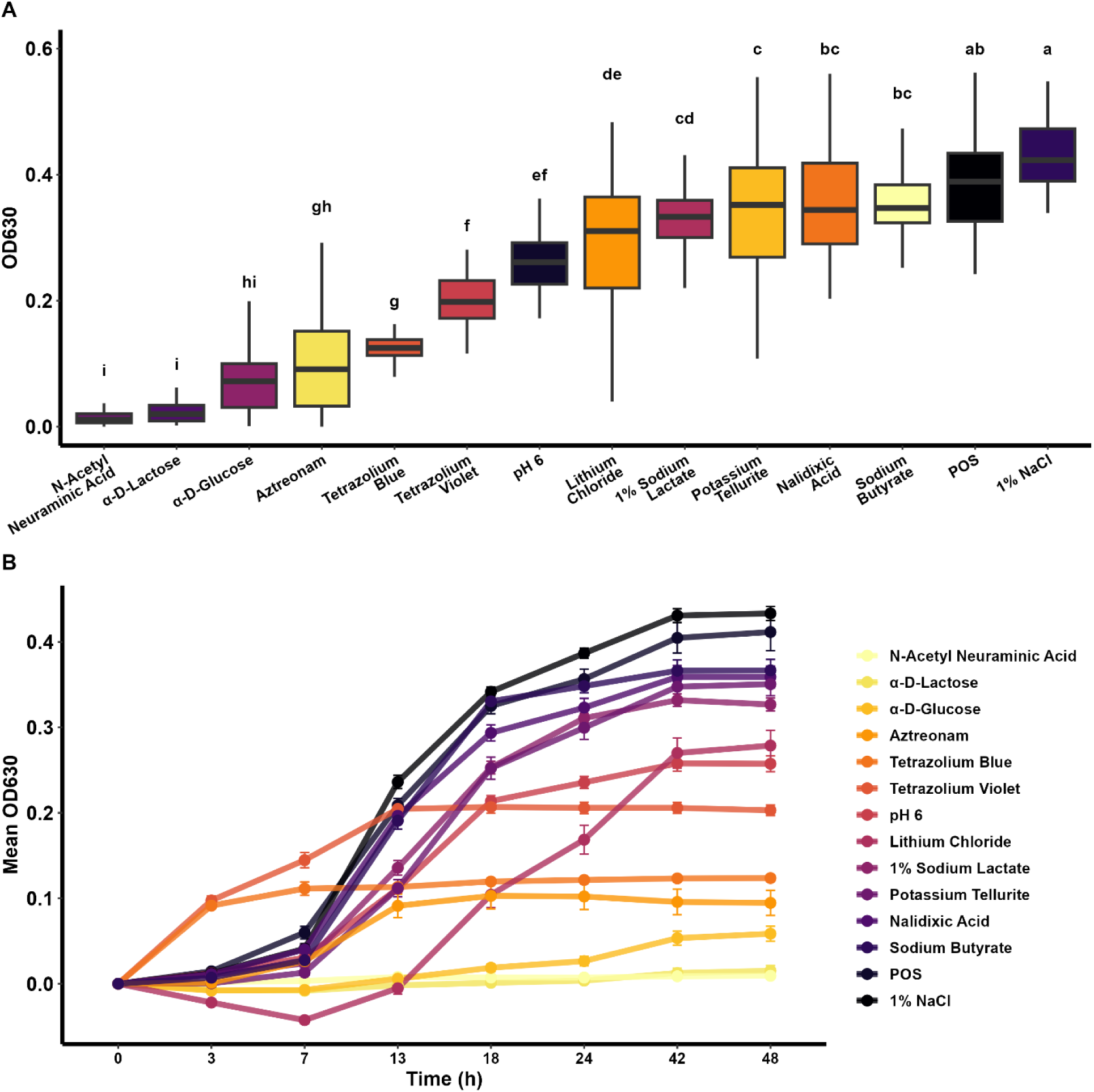
Metabolic characterization of *T. pyogenes* isolates showing A) the preferred substrates in average utilized in order from least to most metabolic activity, and B) their metabolization over time measured by the color change in the Gen III Microplate (n = 49). Different letters indicate different mean OD values (α = 0.05), POS = Positive control well.

**Fig. 9.**
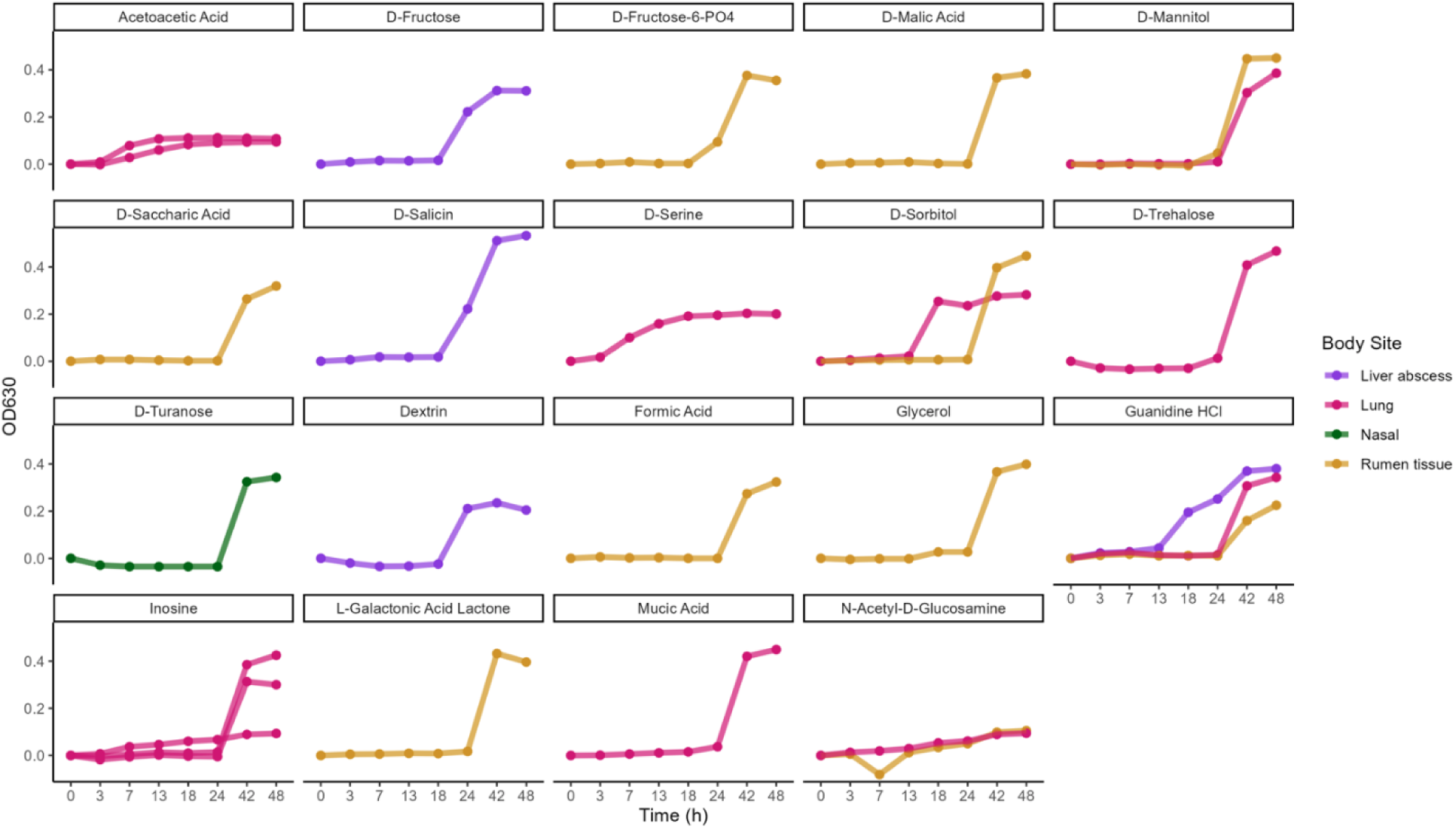
The growth curve of *T. pyogenes* isolates with a longer lag phase based on the metabolism of different substrates.

## Discussion

*Trueperella pyogenes* is an opportunistic pathogen causing suppurative infections, such as mastitis, metritis, pneumonia, and abscesses in livestock, pets, and wildlife, leading to significant health and economic losses. Zoonotic infections of *T. pyogenes* have been reported in humans (Deliwala et al., 2020; Gahrn-Hansen & Frederiksen, 1992; Levy et al., 2009; Plamondon et al., 2007; Stuby et al., 2023). Virulence factors and AMR may contribute to pathogenesis in multiple host species and tissues and increase the risk of ARG dissemination and treatment failure. In the present study, we evaluated the genomes, pangenome, virulence genes, and ARGs of *T. pyogenes* isolated from diverse animal hosts (11 different hosts) comprised of livestock, companion, and wild animals, and from 16 different body sites.

A phylogenetic tree of 83 *T. pyogenes* genomes revealed that the bovine and swine isolates largely clustered into separate clades with the exception of a few bovine isolates that were closely related to the swine isolates. However, we did not observe any grouping based on type of infection or body site between or within animal species. Our analysis revealed that *T. pyogenes* has an open pangenome, similar to what was observed in a previous pangenome analysis of 20 *T. pyogenes* genomes (Thakur et al., 2022). Although other studies have characterized *T. pyogenes* isolated in China, India, and Europe (Hijazin et al., 2011; Moreno et al., 2017; Thakur et al., 2022), genomic characterization of clinical *T. pyogenes* isolates from North America has not previously been done.

The predominant virulence factor genotype was *plo*, *fimA*, *fimC*, *fimE,* and *nanP* (genotype VIII) and *plo, fimA, fimE*, and *nanP* (genotype XVI) which were both found in 14.5% of isolates. All but one *T. pyogenes* genome carried the *plo* gene, and 97.6% of strains from this study also encoded *fimA*. There is great variation in virulence genes distribution among studies evaluating *T. pyogenes*, with some harboring all known virulence genes as the predominant genotype (Ashrafi Tamai et al., 2021), highlighting the versatility of this bacteria as an opportunistic pathogen. Other studies have also identified the *plo* and *fimA* genes in all *T. pyogenes* isolates from bovine mastitis and metritis samples (Rezanejad et al., 2019), similar to our bovine strains (100% carried *plo* and *fimA*). In the present study, two of the seven known *T. pyogenes* virulence genes screened, *plo* and *fimA,* were present in nearly all isolates. This is consistent with other studies of virulence genes of *T. pyogenes* (Ashrafi Tamai et al., 2018; Risseti et al., 2017; Rzewuska et al., 2012, 2016; Silva et al., 2008). The *plo* gene encodes for a hemolytic exotoxin, the cholesterol-dependent cytolysin *plo*, belonging to the family of cholesterol binding molecules in the host cell lipid membrane. Gram-positive pathogens that express these pore-forming toxins are associated with higher cholesterol concentrations in the target cell membranes (Evans et al., 2020). The *plo* protein binds to host cell membranes, inducing the formation of oligomeric β-barrel pores, creating channels (Morton et al., 2019) and leading to cell lysis, in addition to the hemolysis, immune cell lysis, and cytokine expression by the host immune system (Jost & Billington, 2005). A previous study has demonstrated that *plo* is necessary for hemolysis through the observation of non-virulent *T. pyogenes plo* knockout mutants that were not capable of hemolysis when injected into mice (Billington et al., 1997). Given that *plo* and *fimA* were present in almost all 83 *T. pyogenes* genomes analyzed, these two genes may be good targets for vaccine development.

Neuraminidases, encoded by *nanH* and *nanP* in *T. pyogenes*, are important virulence factors for bacteria that inhabit the mucous layer of organs and nutrient-limited environments (Risseti et al., 2017). These enzymes remove sialic acid from polysaccharides, glycolipids, and glycoproteins, which in turn are used as a carbon source by the bacterium, facilitating tissue colonization by reducing the mucous layer viscosity (Byers et al., 1997; Godoy et al., 1993; Tong et al., 2000). Neuraminidases are important in *T. pyogenes* cell adhesion (Jost et al., 2001, 2002) as they are collagen-binding proteins encoded by *cbpA* (Esmay et al., 2003) and fimbriae (Machado et al., 2014; Santos et al., 2010). The high prevalence of the *fimC* and *fimE* genes among *T. pyogenes* isolates recovered from clinical mastitis cases (Zastempowska & Lassa, 2012) suggests that they may be important in mammary gland colonization and maintenance in the uterus of cows. Risk of *T. pyogenes* mastitis is higher at early lactation and increases with reoccurring infections (Duse et al., 2021). In another study, *T. pyogenes* from livestock animal samples demonstrated a difference in the prevalence of *cbpA* between animal hosts, where 100% of reptiles, 25% companion animals, 54% pigs, 7% bovine, 0% horse, rabbit, and rat isolates carried the gene (Hijazin et al., 2011). However, previous studies failed to determine the association between virulence genes and animal host or infection site (Ashrafi Tamai et al., 2021; Hijazin et al., 2011; Risseti et al., 2017; Rzewuska et al., 2012).

*Trueperella pyogenes* is a primary pathogen for uterine infection, and purulent uterine disease in cows has been associated with endometrial cell sensitivity to the *plo* exotoxin causing hemolysis and cytolysis and an inflammatory response with increase of IL-1beta, IL-6, and IL-8 accumulation (Amos et al., 2014). Additionally, the severity of metritis clinical symptoms was linked to the virulence genotype of *T. pyogenes* in postpartum cows (Ashrafi Tamai et al., 2021). This interaction with the animal host immune system can help explain the opportunistic nature of *T. pyogenes*. For instance, about 40% of cows can develop pyometra after parturition, which is linked to the loss of the epithelial cell layers that can provide protection against invasion by *T. pyogenes, E. coli*, and other pathogens (Machado et al., 2014). In addition to virulence factors, *T. pyogenes* can survive inside host phagocytes, shown to persist inside macrophages for 72h (Jost & Billington, 2005) and is capable of forming biofilms that significantly increase their ability to evade the host immune response, attach to epithelial cells, and survive exposure to antibiotics (K. Zhao et al., 2013). Biofilm formation is usually linked to the persistence of infections such as bovine mastitis caused by *Staphylococcus aureus* due to the production of biofilm-associated protein (*Bap*) (Cucarella et al., 2004). Resistant to antibiotics as a result of biofilm formation can possibly explain why treatment for *T. pyogenes* often takes a prolonged period of time for success.

No single ARG was found in all genomes, and the most common ARGs found in these isolates are those confer resistance against tetracyclines, glycopeptides, and MLS_B_. Genotypic multidrug resistance (ARGs to 3 or more antimicrobial classes) was found in 43.3% (36/83) of *T. pyogenes* isolates evaluated in the present study. Isolates that harbored the largest number of ARGs (n = 5-8 genes) were originated from the bovine ruminal tissue (n = 6 isolates), bovine lung (n = 2), swine lung (n = 1), and white-tailed deer lung (n = 1). The isolates without any identified ARGs were originated from the lungs of bovine, swine, and white-tailed deer samples, feline nasal swab, and bovine rumen tissue samples. *T. pyogenes* isolates from this study had ARGs predominantly against tetracyclines, followed by MLS_B_, and glycopeptides. This observation is consistent with other studies evaluating the genotypical AMR profiles of *T. pyogenes*. For example, Werckenthin et al. (2007) observed a high prevalence of resistance against tetracycline and sulfonamides in bovine and swine associated *T. pyogenes* isolates.

Bovine ruminal tissue isolates *T. pyogene*s 2022_1_29 and 2022_1_77 both carried the same eight ARGs conferring resistance to six different antimicrobial classes and appeared to be from the same strain based on ANI (Table S2). Notably, four of these genes *sul1*, *qacEdelta1*, *aadA2*, and *ant(2’’)-Ia* were co-located together on a contig that was potentially from a plasmid. The best matches (>99.5% identity and 90% coverage) for this contig in the NCBI nucleotide database were to species in the *Enterobacterales* order such as *E. coli* (CP050163.1)*, Citrobacter freundii* (CP038659.1)*, Proteus mirabilis* (CP137083.1), and *Salmonella enterica* subsp. *enterica* (OX442404.1). This same contig was also identified in *T. pyogenes* 2022_1_75. A different cattle ruminal tissue-assocated isolate, 2023_3_67, also encoded eight ARGs conferring resistance to five different antimicrobial classes. In this isolate, the *erm*(X) and *tet*(33) genes were co-located on a 8,048 bp contig that was nearly identical (99.95% identity and 100% coverage) to a plasmid (pAP2; AY255627.1) in *T. pyogenes* (Jost et al., 2003). The *erm*(X) and *tet*(33) genes were also found on a plasmid-like contig in isolates 2022_1_30 and 2022_1_96; however, in strains 2022_1_27 and 306 they appeared to be chromosomally encoded as they were located on contigs that were approximately 1.3 Mbp in length (data not shown). Some infections that have *T*. *pyogenes* as one of the primary causative agents, such as endometritis in dairy cows, are treated with antimicrobials. If AMR increases among *T. pyogenes* or spreads between *T. pyogenes* and the other bacterial species through horizontal gene transfer this can lead to a reduction in treatment efficacy and economic losses due to lower milk yields and decreased fertility in the herd. Choosing the correct antimicrobial based on the intrinsic and acquired AMR of *T. pyogenes* is therefore important.

Unfortunately, there are no established breakpoints for *T. pyogenes* based on the disk diffusion assay. Therefore, resistance breakpoints for *S. pneumoniae* and *Corynebacterium* spp. and related coryneform bacteria were used in the present study. Therefore, caution is warranted when extrapolating our results to clinical settings. The *in vitro* disk diffusion assay demonstrated that *T. pyogenes* isolates from the present study were predominantly resistant to tetracycline (75%), clindamycin (37.5%), and erythromycin (24.5%), with a smaller percentage resistant to gentamicin (6.1%) and chloramphenicol (2.1%). Conversely, our isolates were completely susceptible to penicillin, sulfamethoxazole/trimethoprim, and vancomycin. This is in accordance with other studies, where resistance to tetracycline, as high as 85.5% of tested isolates (Zastempowska & Lassa, 2012), and macrolides, (around 30-35% of isolates) (Ashrafi Tamai et al., 2018, 2021; Trinh et al., 2002), are predominant.

Some of the T. pyogenes genomes had the co-occurrence of ARGs and MGE genes (integrons). Integrons can move ARGs through site-specific recombination and are associated with the dissemination of ARGs, via chromosome to plasmid, or plasmid to chromosome (intracellular mobility), and between bacteria and other organisms (intercellular mobility) (Partridge et al., 2018). This class 1 integron is found in the chromosome of betaproteobacteria also associated with the qacEdelta1 and sul1 ARGs (Partridge, 2011), highlighting the possible acquisition of ARGs from the microbial community around these isolates.

Differences in *T. pyogenes* antimicrobial susceptibilities *in vitro* can be observed depending on the type of antimicrobials used, sample origin, and animal species. For instance, *T. pyogenes* of bovine origin exhibited resistance to tetracyclines that differed based on the tetracycline (6% doxycycline, 21% tetracycline, and 88% oxytetracycline) (Yapicier et al., 2022), and infection site (11% in metritis vs. 35% in mastitis) (Ashrafi Tamai et al., 2018, 2021). *T. pyogenes* susceptibility can also differ based on host species and often bovine and swine isolates show diminished susceptibility to antibiotics used in animals (i.e. tetracyclines, sulfonamides, and clindamycin) than isolates from other livestock and wildlife species (Kwiecień et al., 2021; Moreno et al., 2017; Rzewuska et al., 2016). This is likely due to the greater direct exposure to antimicrobials in livestock than their wildlife counterparts.

Some antimicrobials, such as tetracycline and macrolides have traditionally been used in subtherapeutic doses in cattle and swine production systems to promote growth and prevent disease (FDA, 2018). *T. pyogenes* infections often involve multiple bacterial species and, consequently, they are not usually the main target for antimicrobial treatment. However, beta-lactams (penicillins, cephapirin, and ceftiofur), tetracyclines, MLS_B_ (erythromycin, tylosin, clindamycin), trimethoprim-sulfamethoxazole, and enrofloxacin are typically indicated for *T. pyogenes*-related infections (Rzewuska et al. 2019). As observed in this study, and with cattle in France (Guérin-Faublée et al., 1993) and Switzerland (Marchionatti et al., 2024), as well as swine in China (Dong et al., 2019), *T. pyogenes* is widely susceptible to beta-lactams, compared to tetracyclines and MLS_B_. The most effective antimicrobials against *T. pyogenes* isolates tested in the present study were ciprofloxacin, sulfamethoxazole/trimethoprim, penicillin, and vancomycin. Known beta-lactamase genes conferring resistance to the beta-lactams were not found in any of the *T. pyogenes* isolates and they were all classified as susceptible based on their phenotype. Comparisons between studies is limited due to differences in methods (microbroth vs. disk diffusion) and the breakpoints used (CLSI M45, other studies).

When assessing the presence or absence of ARGs in *T. pyogenes* isolates with their phenotypic resistance against the respective antibiotics, the ZOI was significantly (*P* < 0.05) smaller when the ARGs were present as expected. Some isolates had a reduced susceptibility to certain antimicrobials even when no known ARGs for that antimicrobial or antimicrobial class were identified, suggesting that resistance may have been due to a novel antimicrobial gene or other mechanism. It has previously been demonstrated that factors such as silent (unexpressed) genes, low copy numbers, and/or a large distance from the promoter region, can lead to a mismatch between the genotypic and phenotypic antimicrobial susceptibility profiles (Enne et al., 2006; Moran et al., 2017; Sunde & Norström, 2005). The expression of phenotypic resistance without a corresponding ARG in the genome indicate that other mechanisms may play a role in conferring resistance to that antibiotic, including multidrug resistance mechanisms, such as efflux pumps (Piddock, 2006).

The greatest difference in ZOI diameters was observed for tetracycline, erythromycin, and clindamycin, especially between swine and bovine isolates. Cross-resistance to antimicrobials in the MLS_B_ class is expected in isolates carrying the *erm*(X) gene due to their structural similarities (Pernodet et al., 1996). Other mastitis pathogens are also known to carry MLS_B_ resistance genes, highlighting the need for judicious use of those antimicrobials for therapeutic purposes (Li et al., 2015). Isolates from the majority bovine clade (Fig. 1) showed greater resistance to clindamycin, erythromycin, and tetracycline when compared to isolates in the swine clade, indicating possible host-specific AMR differences. Globally, the US ranks as the third largest user of veterinary antimicrobials by volume and it is projected that this will increase 3.8% by 2030 (Tiseo et al., 2020). Antibiotics are often used in livestock for one of three goals: growth promotion, prevention or metaphylaxis, and therapeutic. The most commonly used antibiotics in this industry are tetracyclines, penicillins, macrolides, sulfonamides, aminoglycosides, lincosamides, cephalosporins, and fluroquinolones (FDA, 2018). However more comprehensive analysis of the distribution and correlation between drugs used in animal production and genotypical/phenotypical AMR is needed.

Horizontal gene transfer in the gut can spread ARGs among commensal and pathogenic bacteria (Hayes, 2001; Huddleston, 2014). The spread of AMR in the microbiome of animals as well as farm workers has been demonstrated through horizontal gene transfer via plasmids, with tetracycline medicated feed given to poultry (Levy Stuart B. et al., 1976), nalidixic acid in bovine and swine (Marshall et al., 1990), tetracycline and sulfonamide resistance gene-carrying plasmids in bovine (Oppegaard et al., 2001), and methicillin-resistant *S. aureus* from farm animals (Feßler et al., 2012). Increase in food/meat demand and consequent intensification of farming practices for livestock are associated with the increase in genotypical and phenotypical prevalence of AMR in animals (Wang et al., 2023). This has the potential to also impact AMR in humans (T. P. Van Boeckel et al., 2017; van Bunnik & Woolhouse, 2017) as we see a trend of increased AMR in chickens, pigs, and cattle observed in mid- and lower-income countries for the past two decades (T. Van Boeckel et al., 2019). Factors potentially contributing to increased AMR in livestock include inappropriate and ineffective initial treatment, delaying success in combating the infection, and widespread use (Simoneit et al., 2015). Other studies have observed that human origin *T. pyogenes* isolates are genetically close to bovine isolates (Marchionatti et al., 2024). Spillover from animal hosts to humans who are less adapted to the infection can be susceptible to multidrug resistant strains and treatment is high risk of infection.

Despite our contribution of 59 new *T. pyogenes* genomes in the present study, there are still not many publicly available *T. pyogenes* genomes, especially those from animal species other than cattle and pigs. As *T. pyogenes* has been isolated from multiple domestic and wild animal species, more isolates are needed from these underrepresented sources to better understand host-specific differences, if any, for AMR and virulence of *T. pyogenes*. This can assist in understanding how this bacterial species is well adapted to various microbial niches and can cause infections in different tissues in multiple animal species while remaining a commensal member of the microbiota. The characterization of the *T. pyogenes* genomes revealed variability in virulence gene and ARG profiles. This characterization can assist in molecular epidemiological and evolutionary studies, helping to determine the virulence and AMR distribution and contribution between different animals host species. Although there are no validated breakpoints for *T. pyogenes*, currently, determining genotypic and phenotypic AMR can support antimicrobial therapy choices for this pathogen in clinical settings, a crucial step in the prevention of the development and horizontal spread of ARGs among animal and zoonotic pathogens. Evolution and spread of AMR from farm animals is important because it leads to the failure of treatment of infectious diseases due to drug-resistant bacteria in animals and humans, decreased production, and worsen health and welfare conditions for livestock. The association between genotypical and phenotypical antibiotic resistance highlights the importance of ARGs transfer among pathogenic and zoonotic bacteria, through horizontal transfer of other mechanisms to public health.

The preferred carbon source utilization, in addition to pH and salinity tolerance were similar among all tested *T. pyogenes* isolates (Fig, 9). The OD_630_ was higher for 1% NaCl than the positive controls, suggesting that this osmotic concentration is favorable to the growth of *T. pyogenes*, similarly lactic acid bacteria growth is also enhanced by the addition of 1% NaCl to medium (Korkeala et al., 1992). *T. pyogenes* isolates in general metabolize sodium butyrate, a microbial short-chain fatty acid known for its pro-apoptotic effects. Derived from glucose metabolism, sodium butyrate serves as a nutrient source that supports bacterial growth and potentially influences host-pathogen interactions, it has bactericidal effect against Gram-negative bacteria by inhibiting DNA synthesis, its optimal bactericidal activities occur in low doses, in higher doses bacteria can survive (Crumplin & Smith, 1975). Furthermore, *T. pyogenes* grows in the presence of potassium tellurite, which is used in media selective for *Corynebacterium* species, a compound toxic to many bacteria but inherently tolerated by certain Gram-positive organisms, such as *Corynebacterium diphtheriae* and *S. aureus* (dos Santos et al., 2015; Franks et al., 2014). This resistance extends beyond telluride to encompass other antimicrobials and oxidative stressors, conferring a competitive advantage in hostile environments. Additionally, *T. pyogenes* metabolizes 1% sodium lactate, commonly used as a food preservative due to its pH-lowering and water activity-reducing properties (White et al., 2022). Moreover, the ability of *T. pyogenes* isolates to degrade lithium chloride, albeit rare, suggests a broader spectrum of metabolic versatility potentially mediated by specific genetic mechanisms such as cation transporters. This underscores its adaptability to nutrient-depleted environments, where it can utilize alternative carbon sources to sustain growth and survival.

Conversely, there were some exceptions to the core metabolizable carbon sources and resistance to chemical compounds. A few isolates (Table S5) demonstrated the ability to metabolize diverse carbon sources and grow in the presence of chemical compounds that the remaining majority of the *T. pyogenes* isolates could not. This metabolic flexibility may contribute to their survival, offer a competitive advantage, and a pathogenic potential to them. One advantage lies in their capability to utilize complex carbohydrates like D-trehalose and D-fructose-6-PO4, which serve as an energy source and offer protection environmental stresses, such as osmotic stress and freezing (Nørregaard-Madsen et al., 1995).

The *T. pyogenes* isolate capable of metabolizing D-fructose-PO4 as a sole carbon source came from bovine ruminal tissue. Commensal bacteria in the rumen can travel to the liver and cause liver abscesses. Certain species as *Fusobacterium necrophorum* and *Bifidobacterium* spp. found to be resistant to bile were also determined to have high phosphoketolase activity when compared to their susceptible counterparts. This enzyme is responsible for breaking down fructose-6-PO4 during carbohydrate metabolism (Sánchez et al., 2004), therefore resistance to bile is an advantageous adaptation. Moreover, metabolic pathways include the utilization of organic acids, such as L-Galactonic acid, d-malic acid, mucic acid, saccharic acid, acetoacetic acid, and formic acid, which have bactericidal properties (Entani et al., 1998; Rudrappa et al., 2008). A ruminal tissue *T. pyogenes* isolate could metabolize formic acid, this is an one-carbon product of ruminal acetate fermentation, it represents about 5% of volatile fatty acids in the rumen (Hook et al., 2010). By metabolizing these acids, *T. pyogenes* potentially overcomes environmental pressure and gains a competitive edge in hostile environments, such as ruminal tissues where formic acid is abundant.

Furthermore, *T. pyogenes* utilized sugars like D-sorbitol and D-mannitol, which are prevalent in plant tissues and host environments. D-Mannitol is metabolized by the mannitol-1-phosphate dehydrogenase enzyme in *S. aureus* (Nguyen et al., 2019), which also helps with pH, and osmotic tolerance (Aldridge et al., 1997; Nissen et al., 2005). The mannitol pathway is a strategy employed by some microorganisms to overcome environmental pressures, by reversibly transforming mannitol-1-phosphate (M1P) into fructose-6-phsphate (F6P) with M1PDH. Simply, mannitol is converted to fructose by the mannitol-2-dehydrogenase (M2DH) enzyme, in organisms such as *S. aureus, Clostridium* spp., *Bacillus* spp., *E. coli,* and *Klebsiella pneumoniae* (Nguyen et al., 2019; Ruijter et al., 2003).

Other carbon sources utilized by a few *T. pyogenes* isolates included D-trehalose, a disaccharide produced by algae and plants. Its metabolism can be associated with abiotic stress due to its protective characteristics against osmotic stress and freezing (Argüelles, 2000; Crowe et al., 1990). A few D-trehalose formation/degradation pathways are present in bacteria (Avonce et al., 2006) and its metabolism was linked to pathogenicity (Tournu et al., 2013). The bovine *T. pyogenes* isolate that was able to use D-trehalose as a sole carbon source came from a lung infection. Pathogens and opportunistic pathogens can utilize mannitol as a carbon source. In addition to carbon sources, *T. pyogenes* exhibits the capacity to metabolize amino acids such as D-serine and nucleosides like inosine. Curiously, one *T. pyogenes* isolate from ruminal tissue was able to metabolize glycerol; this molecule can enter the bacterial cell through the glycerol facilitator GlpF protein and is converted into glycerol-3-phophate by the action of the glycerol kinase (GlpK) and glycerol dehydrogenase enzymes in the cytoplasm (Darbon et al., 1999; Zhu et al., 2002). Other bacterial species that can grow with glycerol as a sole carbon source include *Citrobacter freundii*, *K. pneumoniae*, *Clostridium* spp. *Enterobacter* spp. and *Lactobacillus* spp. (Barbirato & Bories, 1997; Biebl, 2001; da Silva et al., 2009; Ito et al., 2005; Seifert et al., 2001; Talarico et al., 1990).

The concentration and availability of these carbon sources and amino acids vary depending on the site within the host (Anfora et al., 2007; Hallam et al., 2023). Different nutrient availability and bacterial adaptation to the nutrient availability can play a role in niche-specificity (Connolly et al., 2015). Bacteria that have the ability to infect and be pathogenic in diverse host body sites simultaneously carry virulence genes and have their expression influenced by the environmental conditions, as observed in different pathotypes of *E. coli* (Connolly et al., 2015) and virulence capsule expression in *S. pneumoniae* (Troxler et al., 2019). Bacteria face many challenges during host colonization. In particular, the environmental conditions play a pivotal role in the adaptation of commensal and pathogenic bacteria where those that can adapt are able to survive. This metabolic versatility is a critical factor in understanding *T. pyogenes* ecology and its ability to thrive amidst the challenges posed by varying host environments.

Future research should include more isolates, encompassing more animal species and anatomical body sites to create a truly comprehensive database for this pathogen. This would allow for better identification of the factors involved in the development of infection and pathogenesis. In addition, long-read sequencing would enable the full assembly of putative plasmid sequences. The presence of highly conserved genes, like the virulence gene *plo,* could be used as a target for vaccine or therapeutic development. *T. pyogenes* is also often involved in polymicrobial infections; therefore, *in vitro* studies alone may not adequately reveal the *in vivo* pathogenicity and virulence when there are interactions with other bacteria. For example, the formation of liver abscesses in conjunction with *Fusobacterium necrophorum* and *Salmonella spp*. or the interaction between *T. pyogenes* with *S. aureus*, *Streptococcus agalactiae*, and *E. coli* in pathogenesis of mastitis Models involving multiple pathogens need to be tested to evaluate the synergistic effect of the metabolite interactions between said pathogens and their resulting virulence, pathogenicity, and adaptation for infection (Ramsey et al., 2011). Lastly, the zoonotic potential of *T. pyogenes* should not be overlooked and needs to be further investigated.

## Conclusions

The genomic characterization of *T. pyogenes* from different animal hosts and different anatomical body sites revealed some clustering between bovine and swine isolates, without distinction for other species or anatomical body sites. Antimicrobial susceptibility testing revealed that *T. pyogenes* isolates were least susceptible to tetracycline, clindamycin and erythromycin, Overall, bovine isolates displayed decreased antimicrobial susceptibility to tetracycline, erythromycin, clindamycin, ciprofloxacin, and chloramphenicol than that of the other animal hosts. The results of this study suggest that *T. pyogenes* strains are not specific to a particular host or body site. Overall, *T. pyogenes*’ metabolic flexibility and tolerance to environmental stresses, coupled with its capacity to utilize a spectrum of carbon sources and withstand toxic compounds, underscore its adaptive strategies for survival and pathogenicity. These attributes highlight the bacterium’s ability to thrive in challenging environments and contribute to its role as a significant pathogen in various host species.

## Materials and Methods

### Isolate collection and culturing conditions

A total of 60 *T. pyogenes* isolates were recovered from 6 different animal hosts and 11 body sites (Table S1) and subjected to whole genome sequencing. These isolates were from animals originating from the Midwestern United States and isolated by three different labs (n = 27, Diagnostic Medicine/Pathobiology, Kansas State University, Manhattan, KS, USA; n = 30, Veterinary Diagnostic Laboratory [VDL], North Dakota State University [NDSU]; n = 2, Department of Microbiological Sciences, NDSU). Isolates from Kansas State University were isolated using sheep blood agar (Remel, Thermo Fisher Scientific Inc., Lenexa, KS) incubated in a 5% CO_2_ incubator and then stored in brain heart infusion broth (BHI; BD, Franklin Lakes, NJ, USA) with 20% glycerol at −80°C before being shipped to our lab at NDSU. All *T. pyogenes* strains isolated by the VDL were clinical isolates from samples submitted for diagnosis. The remaining *T. pyogenes* strains (93CBB, and 51CBC) were isolated in our laboratory from vaginal swabs taken from healthy beef cattle that were housed and raised at NDSU. In addition, *T. pyogenes* strain NCTC 5224 (ATCC-19411, American Type Culture Collection, Manassas, VA, USA), of swine origin, was included as a reference strain. *T. pyogenes* was isolated on tryptic soy agar supplemented with 5% defibrinated sheep’s blood (TSAb) (BD, Franklin Lakes, NJ, USA) incubated at 37°C with 5% CO_2_ for 24-48 h. Colonies were presumptively identified as *T. pyogenes* based on phenotypic characteristics and confirmed with matrix-assisted laser desorption/ionization time-of-flight (MALDI-TOF) (Olson et al., 2022). Confirmed *T. pyogenes* isolates were cryopreserved in BHI broth with 20% glycerol and stored at −80°C. For genomic DNA extraction and antimicrobial susceptibility testing, cryopreserved bacterial glycerol stocks were re-streaked onto TSAb plates and incubated at 37°C with 5% CO_2_ for 24-48 h.

### Genomic DNA extraction, library preparation, and whole genome sequencing

Isolates (n = 60) were cultured on TSAb overnight at 37°C in 5% CO_2_, and a single colony was then selected, re-streaked onto TSAb and cultured overnight at 37°C in 5% CO_2_ to ensure the purity of the bacterial colony. One colony was used to inoculate 5 ml of BHI broth incubated overnight at 37°C in 5% CO_2_. One ml of overnight culture was subjected to genomic DNA extraction using a DNeasy Blood and Tissue kit (Qiagen, Valencia, CA, USA) with an enzymatic lysis pre-step, as described previously (Amat et al., 2019, 2021). The DNA quality was determined by a NanoDrop ND-1000 spectrophotometer and the concentration measured using the Quant-iT PicoGreen dsDNA kit with a Qubit 4 fluorometer (Thermo Fischer Scientific, Waltham, CA, USA) and then stored at −20°C until library preparation. DNA libraries were constructed using the DNBSEQ short-read library preparation protocols. Libraries were sequenced using the DNA-based nanoball technology DNBSEQ on a DNBSEQ-G50 platform (MGI Tech, Shenzhen, China) with a PE150 flow cell and 2×150 bp sequencing length. After sequencing, raw reads were quality filtered with sequences shorter than 150 bp and >1% N content removed as well as adapter sequences, and low-quality reads (Q < 20) using the SOAPnuke tool (Chen et al., 2018).

### Publicly available T. pyogenes genome acquisition and isolate origins

In addition to the 60 *T. pyogenes* genomes sequenced in the current study, all publicly available *T. pyogenes* genomes in the National Center for Biotechnology Information (NCBI) Genome database were downloaded (n = 22; November 22, 2023) and included in the genomic analyses (Table S1). Only isolate genomes (i.e., non-metagenome-assembled genomes) were included. These additional genomes were included to expand the number of animal hosts and body sites available for analysis to gain insight into the host and niche-specificity. Three *T. pyogenes* ATCC 1941 assemblies were excluded as this strain was also sequenced in the present study.

### Genome assembly and annotation

Default parameters were used for all software unless specified otherwise. Genomes were assembled using SPAdes v. 3.15.5 (Prjibelski et al., 2020) with the “isolate” flag. Assemblies were filtered with BBMap v. 38.96 (https://sourceforge.net/projects/bbmap/) to include only those contigs that were at least 500 bp long. Genome assembly completeness and contamination was assessed with CheckM2 v. 1.0.1 (Chklovski et al., 2023) and genomes with greater than five percent contamination were excluded from further analysis. The quality of the assemblies was evaluated with QUAST v. 5.0.2 (Gurevich et al., 2013) and any assembly with over 100 contigs was discarded. To improve the quality of some of the assemblies with excessive coverage, Seqtk v. 1.4 (https://github.com/lh3/seqtk) was used to subsample the number of reads in each sample to approximately 100X coverage. The completeness and contamination values along with the number and size of the contigs were used to select the best assembly for each sample. Prokka v. 1.14.6 (Seemann, 2014) was used to annotate genomes with the “genus *Trueperella*” flag and a custom *Trueperella* database, which was created by downloading the annotated *T. pyogenes* reference genome ASM61205v1 assembly (GCF_000612055.1) from the NCBI database.

### Pangenome, antimicrobial resistance and virulence gene analysis

The core genome of all *T. pyogenes* assemblies (n = 82) was determined with Roary v. 3.13.0 (https://sanger-pathogens.github.io/Roary/) (Page et al., 2015) and these genes were then aligned with MAFFT v. 7.475 (Katoh and Standley 2013). RAxML-ng v.1.2.0 (Kozlov et al., 2019) was used to create a phylogenetic tree with bootstrapping and the GTR+G substitution model. The phylogenetic tree was visualized with iTol v.6.8.1 (Letunic & Bork, 2021). The average nucleotide identity (ANI) between all 82 *T. pyogenes* genomes was also determined using fastANI v. 1.3.4 (Jain et al., 2018; Rodriguez-R et al., 2024). Antimicrobial resistance genes were detected in the genome assemblies with RGI v. 6.0.1 and CARD v. 3.2.5 (Alcock et al., 2023). Genomes with more than one ARG found on the same contig were visualized with Proksee (Grant et al., 2023) and genes associated with mobile genetic elements were annotated with mobileOG-db v. 1.1.3 (Brown et al., 2022). A custom database of *T. pyogenes* virulence genes was created and included *cbpA* (WP_025296864.1), *fimA* (AHU90603.1), *fimC* (AHU90433.1), *fimE* (AHU90532.1), *nanH* (AAK15462.1), *nanP* (AAK98800.1), and *plo* (WP_025295886.1). All genome assemblies were screened for these genes using DIAMOND v. 2.1.8.162 (Buchfink et al., 2021) with a 90% identity threshold. These virulence genes were chosen because of their reported significance in the pathogenesis of *T. pyogenes*.

### Phenotypic antimicrobial susceptibility testing using disk diffusion

To evaluate phenotypic antimicrobial resistance, a subset of the *T. pyogenes* isolates (n = 49) was selected and subjected to susceptibility testing against 9 different antibiotics using the Kirby-Bauer disk diffusion method (Bauer et al., 1966). These antibiotics were chosen based on the ARGs identified in the genomic analysis. Commercially available dried 6-mm filter paper disks containing specific concentrations of the following antimicrobial drugs were tested: erythromycin (E-15), sulfamethoxazole-trimethoprim (SXT25), tetracycline (Te-30), penicillin (P-10), ciprofloxacin (CIP-5), clindamycin (CC-2), vancomycin (Va-30), gentamicin (GM-10), and chloramphenicol (C-30) (Hardy Diagnostics, Santa Maria, CA, USA). A direct colony suspension of a 0.5 McFarland standard was prepared using 5 mL of sterile 0.85% phosphate-buffered saline (1X PBS, Thermo Fisher Scientific, Waltham, VA, USA) and a colony from a blood agar plate incubated for 24 h at 37°C with 5% CO_2_. Inoculum turbidity was measured using a MicroScan turbidity meter (Beckman Coulter, Brea, CA, USA) and a sterile cotton swab was used to spread the inoculum on the plates. The inoculum was spread in three different directions to ensure an even distribution of the culture and left to dry on the benchtop before disks were applied to the surface of the plate. Plates used for the disk diffusion assay were 100 mm cation-adjusted Mueller-Hinton agar (Oxoid, Basingstoke, Hampshire, UK) supplemented with 5% defibrinated sheep’s blood poured to a minimum depth of 4 mm, as recommended for disk diffusion testing of fastidious organisms. *Streptococcus pneumoniae* (ATCC 49619) was used as an assay quality control (CLSI, 2024) and three disks were used on each agar plate. Disks were removed from their cartridge using a pair of flame-sterilized tweezers, placed onto the agar plate at least 24 mm apart from each other and lightly pressed into the agar. Plates were inverted and incubated at 37°C with 5% CO_2_ for 24 h. After 24 h of incubation, the ZOI was observed and recorded using a caliper to the nearest millimeter. Isolates were classified as resistant, intermediate, or susceptible according to the CLSI-2020 M100 Performance Standards for Antimicrobial Susceptibility breakpoints for *S*. *pneumoniae* (CLSI, 2020) for chloramphenicol, clindamycin, sulfamethoxazole/trimethoprim, tetracycline, vancomycin, and erythromycin, and the coryneform bacteria breakpoints for ciprofloxacin, as breakpoints for *T. pyogenes* are not available and have not been validated to date. Additionally, as no disk diffusion breakpoints are available for either *S*. *pneumoniae,* coryneform bacteria, or *T. pyogenes,* complete resistance (ZOI = 0) was used to classify an isolate as resistant to gentamicin, while any other value for the ZOI was considered susceptible for this study.

### Evaluation of the association between genotypic and phenotypic AMR

The association between the ZOI (in mm) and the presence or absence of ARGs for each of the tested antibiotics was evaluated by the Kruskal-Wallis test as the data was not normally distributed. Normality was tested with the Shapiro-Wilk test and a significance level of α < 0.05 was used. Correlation between the presence of an ARG (genotypic resistance) and antimicrobial susceptibility, resulting from the disk diffusion test, (phenotypic resistance) was determined by Pearson’s Chi-squared test with Yates’ continuity correction, and when the expected values for contingency tables were smaller than 5, then the Fisher’s Exact Test was used. Significance was considered at α < 0.05. All analysis was conducted in RStudio using R v. 4.3.3.

### Metabolic profiling of T. pyogenes isolates

Biology GENIII MicroPlates were used to biochemically profile 49 *T. pyogenes* isolates. Each *T. pyogenes* isolate was tested in 96-well Microplate containing 71 carbon sources and 23 chemical sensitivity analysis (Table S5). The Biolog MicroPlate results show a positive reaction by the change in color to purple due to the reduction reactions of tetrazolium violet present in the substrate. Therefore, the change in color is a result of the bacterial respiration or oxidation of the substrate, rather than cell density from bacterial growth only. It is a method for measuring the adaptation and survivability of the bacteria while exposed to a single carbon source or chemical compound. Measuring the color change over time also has the advantage of showing the differences in how fast bacteria can adapt, if needed, to utilizing a carbon source or how the metabolism is impacted by these different substrates, rather than just looking at the final color change after incubation for a positive/negative reaction.

*T. pyogenes* isolates (Table 1) were transferred from 20% glycerol BHI stocks stored at - 80°C onto TSAb, incubated at 37°C with 5% CO_2_ and sub-streaked twice using the same conditions. From the TSAb plate, a sterile cotton swab was used to transfer the isolated colonies into tubes containing inoculating fluid A (IF-A; Biolog, Hayward, CA) to an OD of 0.2-0.3 using a MicroScan turbidity meter (Beckman Coulter, Brea, CA, USA), according to manufacturer’s instructions. A cell density of 95-92% T, as recommended by the manufacturer for fastidious organisms such as *T. pyogenes,* was used. One hundred microliters of prepared inoculum were transferred into each of the 96 wells of the GenIII plate (Biolog, Hayward, CA) containing 94 biochemical tests, including 71 different carbon sources and 23 chemical sensitivity analysis. Then, the OD_630_ was measured using the Gen5 Microplate reader (BioTek, Winooski, VT, USA) before incubation (0h) and after 3, 7, 13, 18, 24, 43, and 48h of incubation. Differences between the mean final OD values were determined using the non-parametric Kuskal-Wallis test and multiple comparisons adjusted with Dunn Bonferroni method at a significance level α = 0.05, all analysis were done using the RStudio software version 4.3.3.

## Acknowledgments

The authors would like to acknowledge Dr. Kelli Maddock from the Veterinary Diagnostics Laboratory at NDSU for providing *T. pyogenes* isolates and assisting with antimicrobial susceptibility testing and MALDI confirmation of isolates.

## Data availability

All genome assemblies have been deposited in NCBI’s GenBank under the BioProject PRJNA1071155.

## Tables and Figures

**Table S1.**
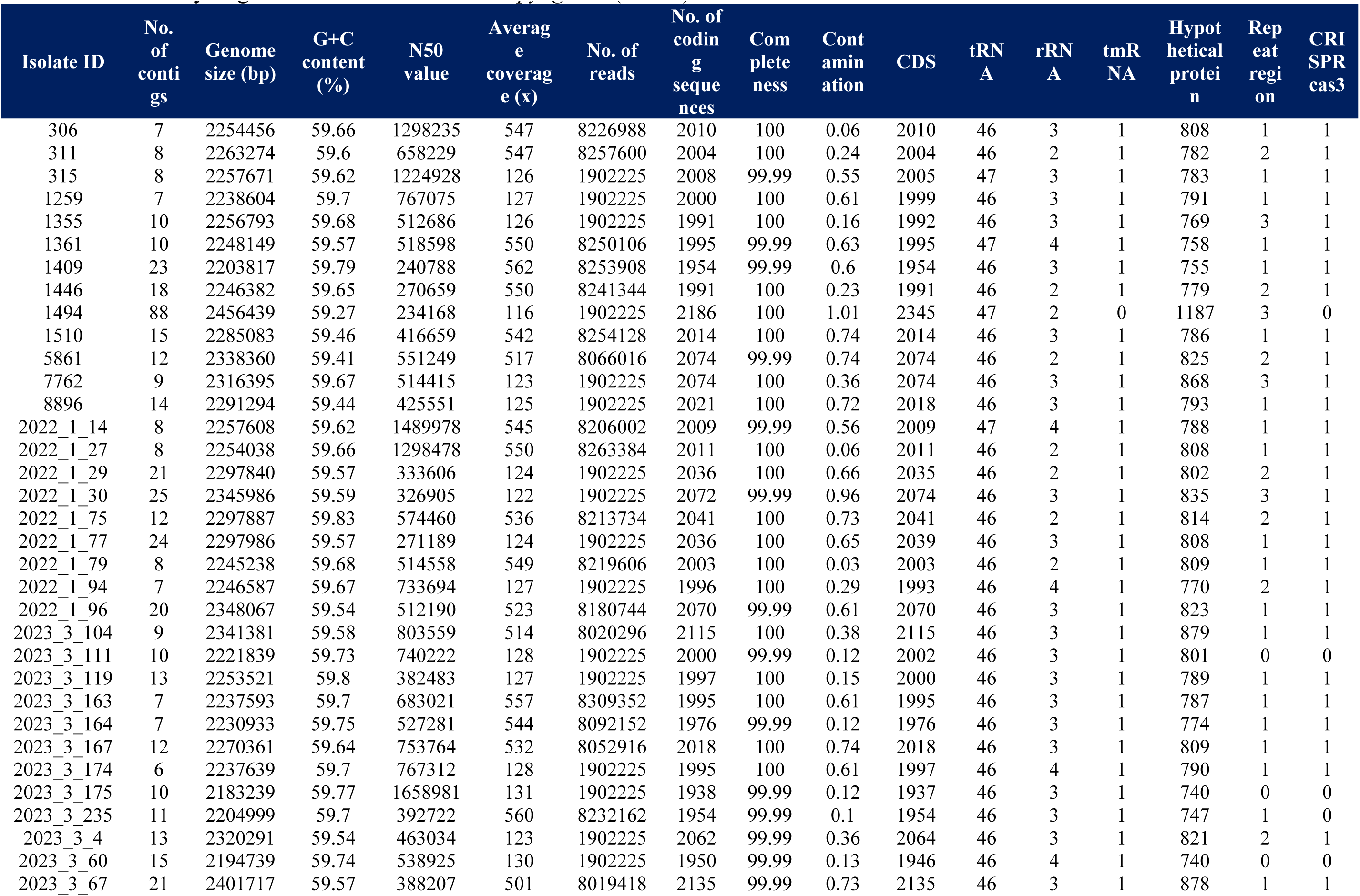
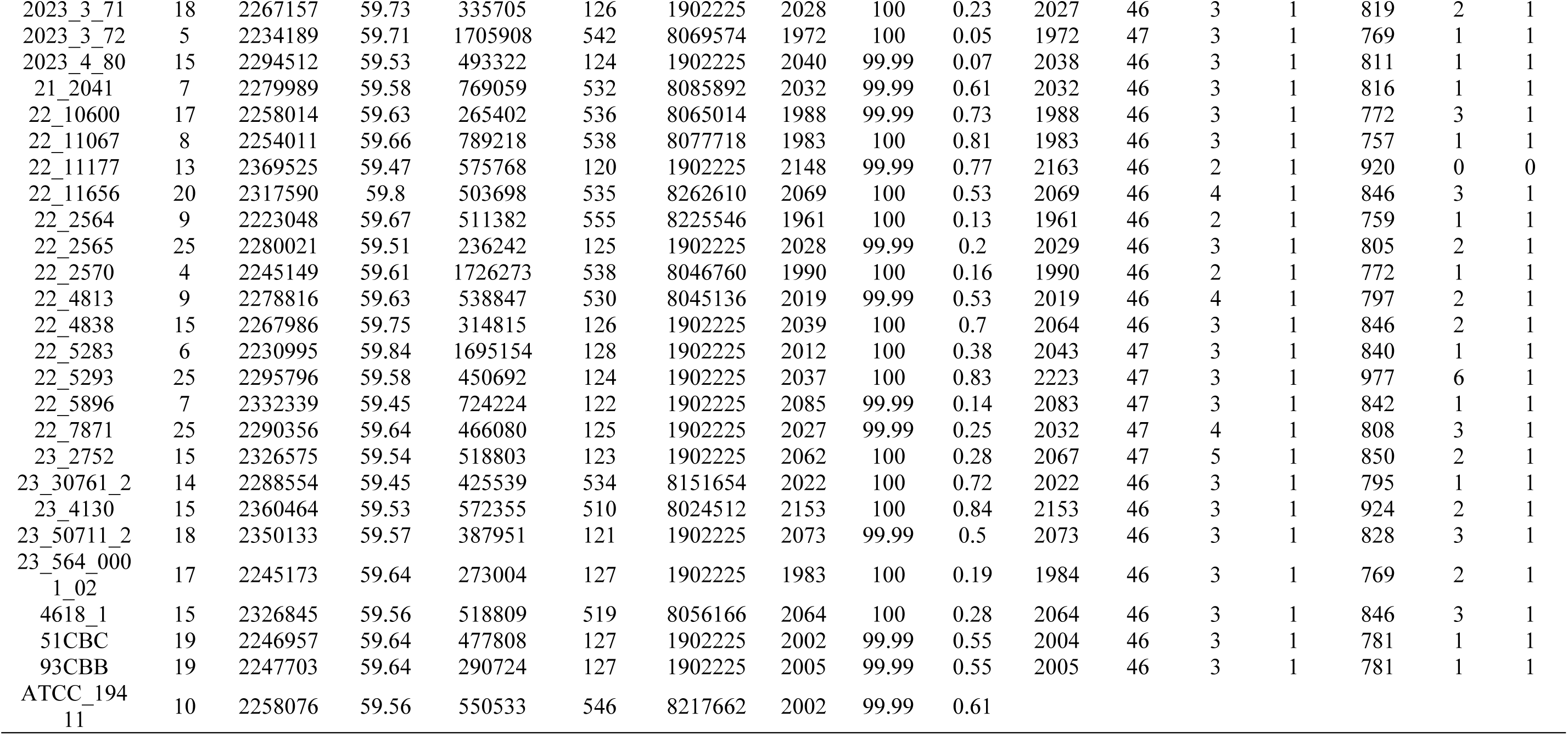
Summary of genomic characteristics of *T. pyogenes* (n = 60)

**Table S2.** Average nucleotide (ANI) of *T. pyogenes* isolates with greater than 99.5% ANI.

| Genome 1 | Genome 2 | ANI | Orthologous<br>fragment<br>count | Total<br>fragment<br>count | ANI % |
| --- | --- | --- | --- | --- | --- |
| 2022_1_14 (bovine) | 315 (bovine) | 100 | 748 | 749 | > 99.99% |
| 2023_3_163 (bovine) | 2023_3_174 (bovine) | 100 | 741 | 742 | > 99.99% |
| 51CBC (bovine) | 93CBB (bovine) | 100 | 736 | 739 | > 99.99% |
| 2022_1_77 (bovine) | 2022_1_29 (bovine) | 100 | 753 | 755 | > 99.99% |
| GCF_003076315 (swine) | GCF_003076295 (swine) | 100 | 790 | 794 | > 99.99% |
| GCF_003076315 (swine) | GCF_003971485 (swine) | 99.999 | 791 | 794 | > 99.99% |
| 2022_1_27 (bovine) | 306 (bovine) | 99.999 | 748 | 748 | > 99.99% |
| GCF_003076295 (swine) | GCF_003971485 (swine) | 99.999 | 792 | 794 | > 99.99% |
| 23_2752 (swine) | 4618_1 (bovine) | 99.996 | 766 | 767 | > 99.99% |
| 1259 (bovine) | 2023_3_174 (bovine) | 99.995 | 741 | 743 | > 99.99% |
| 2023_3_163 (bovine) | 1259 (bovine) | 99.995 | 742 | 742 | > 99.99% |
| 2023_3_111 (bovine) | 2023_3_175 (bovine) | 99.993 | 723 | 736 | > 99.99% |
| 8896 (swine) | 1510 (bovine) | 99.992 | 753 | 756 | > 99.99% |
| 8896 (swine) | 23_30761_2 (deer) | 99.955 | 756 | 756 | 99.5-99.99% |
| 23_30761_2 (deer) | 1510 (bovine) | 99.951 | 753 | 757 | 99.5-99.99% |
| GCF_000612055 (bovine) | GCF_002071825 (bovine) | 99.895 | 668 | 779 | 99.5-99.99% |
| GCF_004009995 (swine) | GCF_004123715 (swine) | 99.848 | 759 | 775 | 99.5-99.99% |
| 2022_1_79 (bovine) | 306 (bovine) | 99.804 | 742 | 744 | 99.5-99.99% |
| 2022_1_27 (bovine) | 2022_1_79 (bovine) | 99.803 | 742 | 748 | 99.5-99.99% |
| 23_50711_2 (bovine) | 2022_1_96 (bovine) | 99.8 | 762 | 775 | 99.5-99.99% |
| 2022_1_96 (bovine) | 2022_1_30 (bovine) | 99.799 | 765 | 773 | 99.5-99.99% |
| 2022_1_96 (bovine) | 2023_3_67 (bovine) | 99.73 | 772 | 773 | 99.5-99.99% |
| GCF_004564055 (swine) | 4618_1 (bovine) | 99.671 | 738 | 757 | 99.5-99.99% |
| GCF_004564055 (swine) | 23_2752 (swine) | 99.667 | 738 | 757 | 99.5-99.99% |
| 23_50711_2 (bovine) | 2022_1_30 (bovine) | 99.636 | 755 | 775 | 99.5-99.99% |
| 1446 (bovine) | 23_564_0001_02 (bovine) | 99.626 | 726 | 739 | 99.5-99.99% |
| 23_50711_2 (bovine) | 2023_3_67 (bovine) | 99.621 | 764 | 775 | 99.5-99.99% |
| 2023_3_67 (bovine) | 2022_1_30 (bovine) | 99.611 | 765 | 791 | 99.5-99.99% |
| 2023_3_60 (bovine) | 2023_3_175 (bovine) | 99.553 | 715 | 723 | 99.5-99.99% |
| 2023_3_60 (bovine) | 2023_3_111 (bovine) | 99.544 | 715 | 723 | 99.5-99.99% |

**Table S3.** Distribution and characteristics *T. pyogenes* genotypes based on the presence of virulence genes (n = 83)

| Genotype | No. genes | Virulence genes | No. genomes | Isolate IDs |
| --- | --- | --- | --- | --- |
| I | 7 | *plo, fimA, fimC, fimE, nanH, nanP, cbpA* | 1 | 1494 |
| II | 6 | *plo, fimA, fimC, fimE, nanH, cbpA* | 1 | UFV1 |
| III | 6 | *plo, fimA, fimC, fimE, nanH, nanP* | 2 | 22_7871, 22_2565 |
| IV | 6 | *plo, fimA, fimC, fimE, nanP, cbpA* | 5 | 2023_3_111, 2023_3_175, 2023_3_4, 2023_3_60, 22/KM0800 |
| V | 6 | *plo, fimA, fimE, nanH, nanP, cbpA* | 1 | 22_2570 |
| VI | 5 | *plo, fimA, fimC, fimE, cbpA* | 6 | 1446, 2023_3_164, 22_4813, 23_564_0001_02, TP6375, Arash114 |
| VII | 5 | *plo, fimA, fimC, fimE, nanH* | 4 | 2022_1_14, 22_5293, 315, EMSSI21 |
| VIII | 5 | *plo, fimA, fimC, fimE, nanP* | 12 | 1355, 1361, 2022_1_75, 2023_3_167, 22_11067, 22_11656, 22_4838, 22_5896, 23_4130, 7762, TP1, TP2 |
| IX | 5 | *plo, fimA, fimE, nanH, cbpA* | 2 | 51CBC, 93CBB |
| X | 5 | *plo, fimA, fimE, nanH, nanP* | 1 | 22_2564 |
| XI | 5 | *plo, fimA, fimE, nanP, cbpA* | 8 | 1409, 2022_1_30, 2022_1_94, 2022_1_96, 2023_3_67, 23_50711_2, 5861, 09KM1269 |
| XII | 4 | *fimA, fimC, fimE, nanP* | 1 | Bu5 |
| XIII | 4 | *plo, fimA, fimC, fimE* | 6 | 2023_3_104, 2023_3_72, 22_5283, 311, EMSSI54, 13KM1326 |
| XIV | 4 | *plo, fimA, fimC, nanP* | 3 | MS249, 22_11177, jx18 |
| XV | 4 | *plo, fimA, fimE, cbpA* | 7 | SH01, 2022_1_29, 2022_1_77, 2023_3_71, 2023_4_80, SH02, SH03 |
| XVI | 4 | *plo, fimA, fimE, nanP* | 12 | 1259, 1510, 2023_3_163, 2023_3_174, 22_10600, 23_2752, 23_30761_2, 4618_1, 8896, TP4479, TP-2849, TP3 |
| XVII | 4 | *plo, fimC, fimE, nanP* | 1 | 2012CQ-ZSH |
| XVIII | 3 | *plo, fimA, fimE* | 5 | 2022_1_27, 2022_1_79, 2023_3_119, 21_2041, 306 |
| XIX | 3 | *plo, fimA, nanP* | 4 | 2023_3_235, TP4, EMSSI48, 13OD0707 |
| XX | 3 | *plo, fimC, nanP* | 1 | ATCC_19411 |

**Table S4.** Zone of inhibition (mm) based on the disk diffusion test for *T. pyogenes* isolates (n = 49) to selected antibiotics.

| Isolate ID | Animal host | Anatomical body site | Chloramphenicol | Clindamycin | Ciprofloxacin | Sulfa/Trimethoprim | Gentamicin | Penicillin | Tetracycline | Vancomycin | Erythromycin |
| --- | --- | --- | --- | --- | --- | --- | --- | --- | --- | --- | --- |
| 306 | Bovine | Liver abscess | . | . | 17 | 34 | 22 | 51 | . | 29 | 23 |
| 311 | Bovine | Liver abscess | 32 | 0 | 16 | 40 | 25 | 53 | 18 | 30 | 0 |
| 315 | Bovine | Liver abscess | 35 | 31 | 16 | 37 | 23 | 52 | 16 | 29 | 44 |
| 1259 | Bovine | Lung | 36 | 34 | 17 | 38 | 26 | 53 | 42 | 32 | 47 |
| 1355 | Ovine | Lung | 33 | 0 | 14 | 36 | 23 | 50 | 11 | 30 | 0 |
| 1409 | Bovine | Lung | 36 | 35 | 22 | 39 | 27 | 52 | 40 | 31 | 49 |
| 1446 | Bovine | Lung | 34 | 0 | 15 | 39 | 26 | 43 | 13 | 32 | 0 |
| 1494 | Deer | Lung | 39 | 36 | 14 | 39 | 26 | 50 | 13 | 29 | 46 |
| 1510 | Bovine | Lung | 39 | 34 | 16 | 43 | 26 | 58 | 45 | 31 | 49 |
| 5861 | Bovine | Lung | 41 | 28 | 16 | 41 | 26 | 57 | 17 | 32 | 43 |
| 7762 | Bovine | Lung | 39 | 35 | 17 | 43 | 29 | 56 | 12 | 33 | 48 |
| 8896 | Swine | Lung | 34 | 31 | 19 | 39 | 24 | 55 | 42 | 32 | 47 |
| 164CBD | Bovine | Vaginal swab | 38 | 34 | 17 | 38 | 25 | 55 | 13 | 31 | 47 |
| 2022_1_29 | Bovine | Rumen Tissue | 33 | 0 | 13 | 36 | 0 | 51 | 11 | 31 | 19 |
| 2022_1_30 | Bovine | Rumen Tissue | 33 | 0 | 14 | 35 | 23 | 47 | 12 | 29 | 0 |
| 2022_1_75 | Bovine | Rumen Tissue | 34 | 0 | 13 | 36 | 0 | 52 | 11 | 30 | 0 |
| 2022_1_77 | Bovine | Rumen Tissue | 36 | 0 | 14 | 33 | 0 | 51 | 11 | 30 | 23 |
| 2022_1_78 | Bovine | Rumen Tissue | 32 | 0 | 16 | 36 | 23 | 51 | 11 | 29 | 0 |
| 2022_1_91 | Bovine | Rumen Tissue | 15 | 0 | 17 | 38 | 25 | 54 | 12 | 32 | 14 |
| 2023_3_167 | Bovine | Rumen Tissue | 37 | 32 | 12 | 37 | 25 | 56 | 39 | 32 | 46 |
| 2023_3_4 | Bovine | Rumen Tissue | 38 | 37 | 14 | 33 | 25 | 51 | 13 | 30 | 46 |
| 2023_3_72 | Bovine | Rumen Tissue | 35 | 32 | 12 | 36 | 22 | 57 | 12 | 31 | 45 |
| 2023_4_80 | Bovine | Rumen Tissue | 34 | 10 | 17 | 40 | 25 | 50 | 12 | 30 | 24 |
| 2023-3-235 | Bovine | Rumen Tissue | 36 | 32 | 13 | 36 | 26 | 49 | 36 | 29 | 45 |
| 21_2041 | Deer | Lung | 40 | 36 | 18 | 40 | 25 | 54 | 13 | 31 | 48 |
| 22_10600 | Feline | Nasal swab | 41 | 34 | 17 | 42 | 25 | 51 | 18 | 33 | 48 |
| 22_11067 | Ovine | Umbilicus | 38 | 33 | 17 | 39 | 26 | 54 | 13 | 33 | 45 |
| 22_11177 | Bovine | Lung | 37 | 33 | 17 | 40 | 25 | 56 | 22 | 31 | 45 |
| 22_11656 | Bovine | Lung | 39 | 0 | 23 | 36 | 26 | 52 | 14 | 31 | 17 |
| 22_2564 | Bovine | Peri fluid | 36 | 33 | 17 | 38 | 27 | 52 | 13 | 31 | 46 |
| 22_2565 | Bison | Lung | 41 | 37 | 17 | 39 | 30 | 53 | 44 | 32 | 47 |
| 22_2570 | Bovine | Milk | 37 | 33 | 18 | 38 | 24 | 54 | 14 | 31 | 45 |
| 22_4838 | Bovine | Lung | 38 | 35 | 18 | 39 | 27 | 54 | 15 | 32 | 47 |
| 22_5283 | Bovine | Lung | 37 | 0 | 15 | 40 | 27 | 59 | 13 | 34 | 0 |
| 22_5293 | Bovine | Foot | 40 | 34 | 18 | 40 | 27 | 58 | 14 | 31 | 46 |
| 22_5896 | Bovine | Lung | 40 | 34 | 19 | 42 | 25 | 57 | 46 | 32 | 49 |
| 22_7871 | Ovine | Mammary | 38 | 34 | 16 | 39 | 25 | 55 | 16 | 31 | 46 |
| 23_2752 | Swine | Lung | 40 | 34 | 19 | 42 | 25 | 60 | 20 | 33 | 53 |
| 23_30761_2 | Deer | Lung | 34 | 28 | 17 | 38 | 23 | 56 | 42 | 30 | 44 |
| 23_4130 | Bovine | Lung | 37 | 0 | 17 | 43 | 28 | 56 | 13 | 32 | 0 |
| 23_564_0001_02 | Bovine | Liver | 31 | 0 | 15 | 39 | 25 | 58 | 11 | 32 | 20 |
| 4618_1 | Bovine | Abscess | 38 | 33 | 19 | 41 | 25 | 51 | 27 | 32 | 47 |
| 51CBC | Bovine | Vaginal swab | 32 | 31 | 17 | 37 | 24 | 53 | 25 | 30 | 44 |
| 93CBB | Bovine | Vaginal swab | 38 | 32 | 20 | 35 | 28 | 51 | 23 | 31 | 53 |
| ATCC_19411 | Swine | . | 38 | 36 | 14 | 38 | 27 | 51 | 46 | 33 | 49 |
| 2023_3_67 | Bovine | Rumen Tissue | 32 | 0 | 16 | 37 | 21 | 45 | 9 | 29 | 22 |
| 2022_1_79 | Bovine | Rumen Tissue | 31 | 0 | 16 | 34 | 20 | 38 | 11 | 28 | 10 |
| 2022_1_96 | Bovine | Rumen Tissue | 37 | 0 | 15 | 35 | 20 | 40 | 10 | 31 | 0 |
| 23_50711_2 | Bovine | Lung | 31 | 0 | 16 | 31 | 17 | 41 | 9 | 30 | 0 |

**Table S5.**
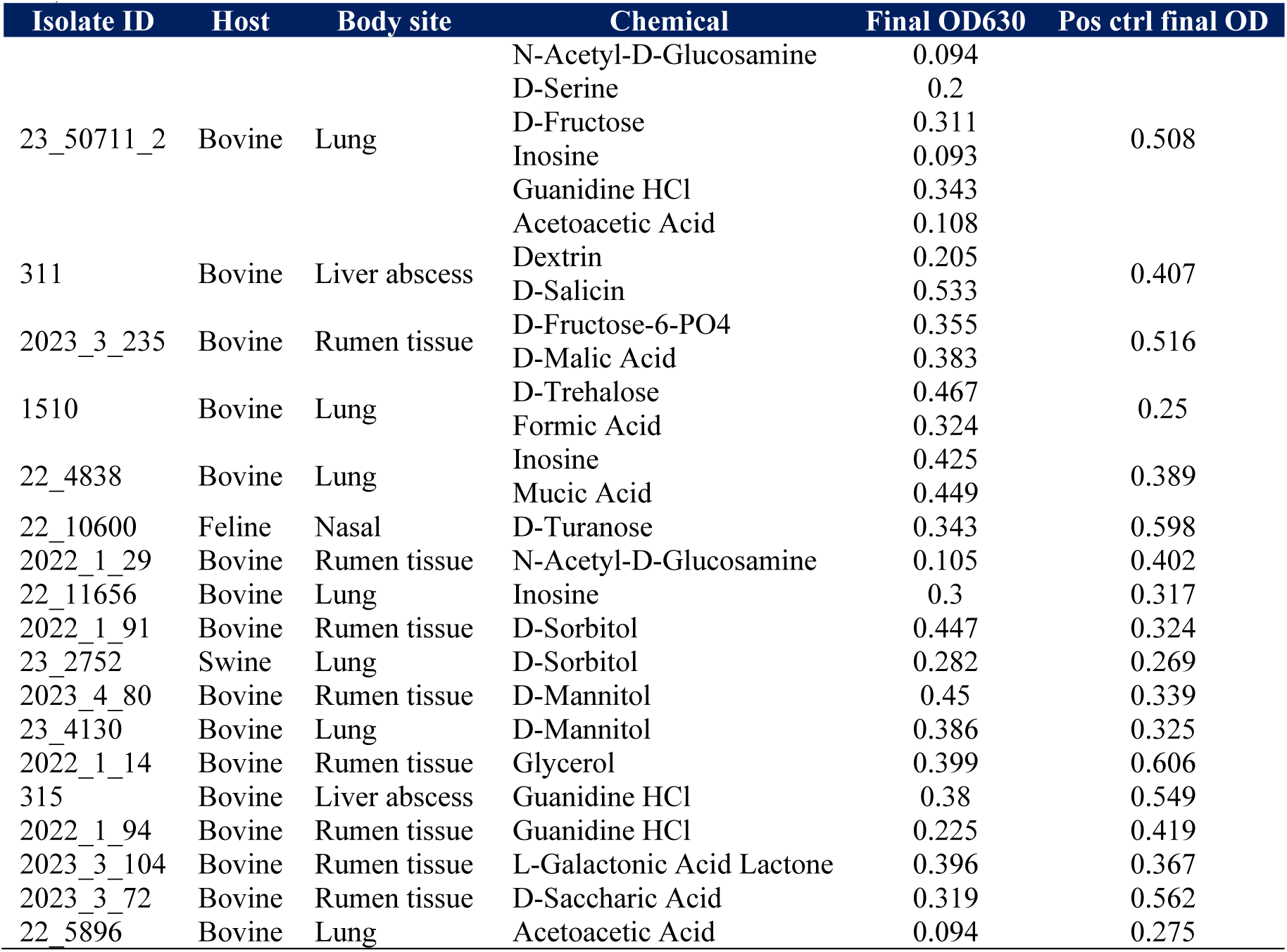
Growth of *T. pyogenes* isolates in different carbon sources and chemical assays (n = 49)

**Figure S1.**
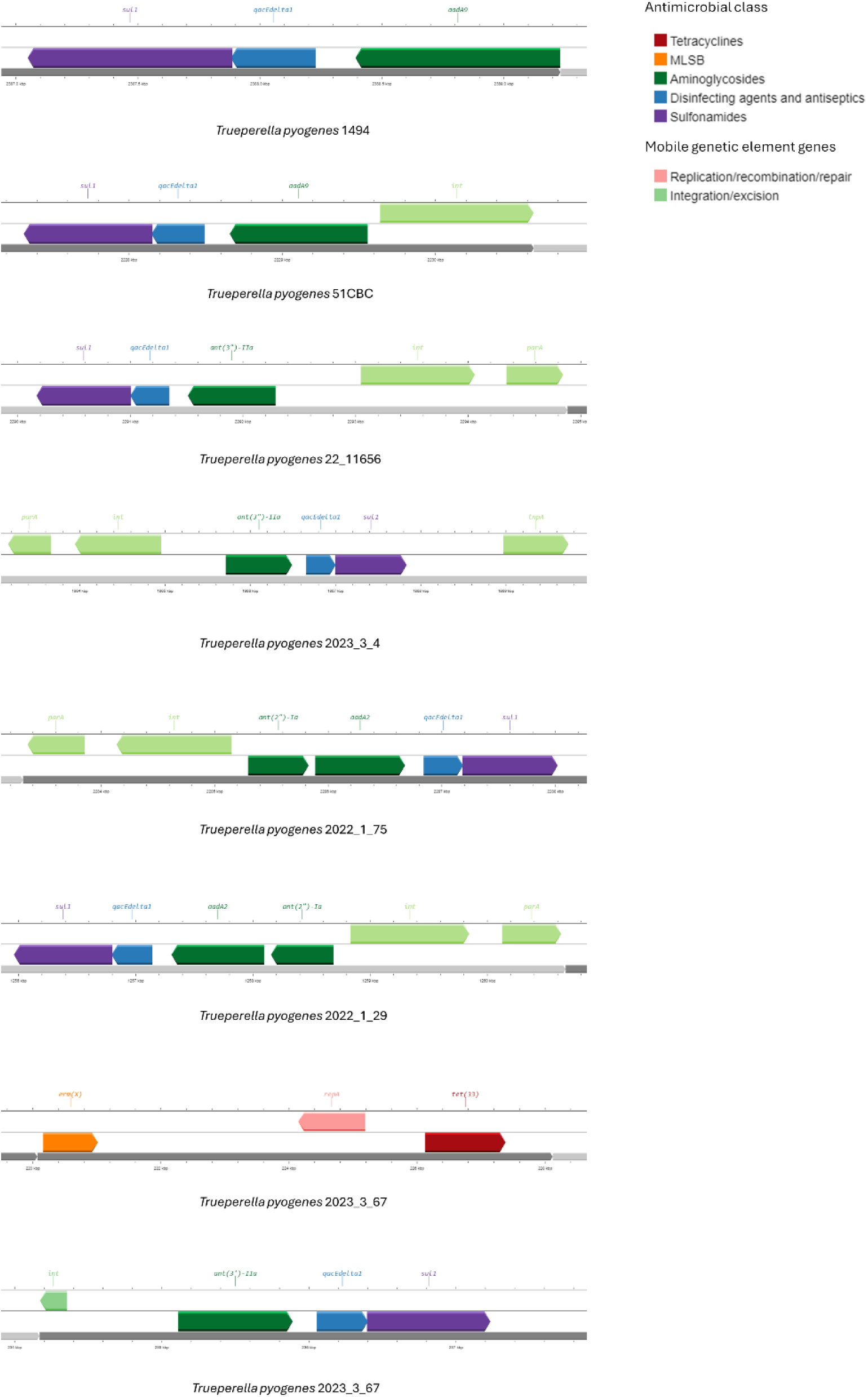
Antimicrobial resistance genes co-located on the same contig in *Trueperella pyogenes* genome assemblies. Also included are mobile genetic element genes.

